# Sexually dimorphic responses to MPTP found in microglia, inflammation and gut microbiota in a progressive monkey model of Parkinson’s disease

**DOI:** 10.1101/2020.01.30.925883

**Authors:** Valerie Joers, Gunasingh Masilamoni, Doty Kempf, Alison R Weiss, Travis Rotterman, Benjamin Murray, Gul Yalcin-Cakmakli, Ronald J Voll, Mark M Goodman, Leonard Howell, Jocelyne Bachevalier, Stefan Green, Ankur Naqib, Maliha Shaikh, Phillip Engen, Ali Keshavarzian, Christopher J Barnum, Jonathon A Nye, Yoland Smith, Malú Gámez Tansey

## Abstract

Inflammation has been linked to the development of nonmotor symptoms in Parkinson’s disease (PD), which greatly impact patients’ quality of life and can often precede motor symptoms. Suitable animal models are critical for our understanding of the mechanisms underlying disease and the associated prodromal disturbances. The neurotoxin 1- methyl-4-phenyl-1,2,3,6-tetrahydropyridine (MPTP)-treated monkey model is commonly seen as a “gold standard” model that closely mimics the clinical motor symptoms and the nigrostriatal dopaminergic loss of PD, however MPTP toxicity extends to other nondopaminergic regions. Yet, there are limited reports monitoring the MPTP-induced progressive central and peripheral inflammation as well as other nonmotor symptoms such as gastrointestinal function and microbiota. The main objective of this study is to gain a broader understanding of central and peripheral inflammatory dysfunction triggered by exposure to a neurotoxicant known to degenerate nigral dopaminergic neurons in order to understand the potential role of inflammation in prodromal/pre-motor features of PD-like degeneration in a progressive non-human primate model of the disease. We measured inflammatory proteins in plasma and CSF and performed [^18^F]FEPPA PET scans to evaluate translocator proteins (TSPO) or microglial activation in a small cohort of rhesus monkeys (n=5) given weekly low doses of MPTP (0.2-0.8 mg/kg, im). Additionally, monkeys were evaluated for working memory and executive function using various behavior tasks and for gastrointestinal hyperpermeability and microbiota composition. Monkeys were also treated with novel TNF inhibitor XPro1595 (10mg/kg, n=3) or vehicle (n=2) every three days starting 11 weeks after the initiation of MPTP to determine whether nonmotor symptoms are tied to TNF signaling and whether XPro1595 would alter inflammation and microglial behavior in a progressive model of PD. Our analyses revealed sex-dependent sensitivity to MPTP that resulted in early microglial activation by PET, acute plasma IL-6 and CSF TNF, and earlier parkinsonism as measured by motor deficits in males compared to female monkeys. Sex differences were also identified in microbiota and their metabolites and targeted short chain fatty acids at both basal levels and in response to MPTP. Both sexes displayed cognitive impairment prior to a significant motor phenotype. Importantly, XPro1595 shifted peripheral and central inflammation, and significantly reduced CD68-immunoreactivity in the colon. As such, our findings revealed a sexually dimorphic inflammatory response to chronic MPTP treatment and suggest that males may have higher vulnerability than females to inflammation-induced degeneration. If these findings reflect potential differences in humans, these sex differences have significant implications for therapeutic development of inflammatory targets in the clinic.

## 1. Introduction

Numerous pre-clinical and epidemiological studies have demonstrated that peripheral and central inflammation accompany and may contribute to the progressive dopaminergic neuronal cell death in Parkinson’s disease (PD) (Gao et al., 2008; McGeer et al., 1988). The existence of ongoing and enhanced inflammatory processes in patients with idiopathic PD is supported by *in vivo* evidence of activated microglia imaged by positron emission tomography (PET) (Gerhard et al., 2006; Ouchi et al., 2009; Terada et al., 2016) and altered inflammatory markers measured in biofluids (Brodacki et al., 2008; Eidson et al., 2017). Moreover, genetic evidence from large genome-wide association studies (GWAS) has identified risk single nucleotide polymorphism (SNP) variants in genes that encode immune function (Hamza et al., 2010; Pierce and Coetzee, 2017) and in genes enriched in distinct subsets of immune cells (Coetzee et al., 2016), further implicating the involvement of the immune system in the pathogenesis of PD. A range of nonmotor symptoms are well recognized to burden PD patients and have been detected years prior to a PD diagnosis (Abbott et al., 2005; Goldman and Postuma, 2014; Ross et al., 2006). It has been hypothesized that inflammation plays a role in the development of PD nonmotor symptoms (Barnum and Tansey, 2012), yet the pathophysiological mechanisms behind the emergence of nonmotor dysfunctions has not been determined. The purpose of this study is to gain a broader understanding of central and peripheral inflammatory dysfunction triggered by chronic exposure to 1-methyl-4-phenyl-1,2,3,6-tetrahydropyridine (MPTP).

The MPTP-treated rhesus macaque monkey model is commonly seen as a “gold standard” model that closely mimics the clinical motor symptoms and the nigrostriatal dopaminergic loss of PD, but the neuropathologic and behavioral effects of the drug extend beyond the ventral midbrain dopaminergic cell groups and parkinsonian motor signs when low doses of MPTP are administered chronically to monkeys and mice (Emborg, 2007; Masilamoni et al., 2011; Masilamoni et al., 2017; Masilamoni and Smith, 2018; Vivacqua et al., 2019) Chronic systemic MPTP administration over several weeks to months allows the slow development of parkinsonism that is more similar to PD, importantly creating a window of time to evaluate early premotor symptoms including inflammation and nonmotor manifestations that are alike impaired in PD. Dysregulated cytokines have been described in non-human primates treated with MPTP (Mondal et al., 2012; Roy et al., 2015) or systemically (Barcia et al., 2005; De Pablos et al., 2009), but the trajectory of the inflammatory environment has not been detailed throughout the gradual progression of MPTP-induced neuropathology and related behavioral changes. In addition to parkinsonian motor signs, chronically MPTP-treated monkeys display early cognitive dysfunction (Roeltgen and Schneider, 1991; Schneider and Kovelowski, 1990; Schneider et al., 2010; Slovin et al., 1999), alterations in sleep, social relationships, olfaction, and gastrointestinal pathologies (Almirall et al., 1999; Barraud et al., 2009; Chaumette et al., 2009; Durand et al., 2015; Phillips et al., 2017). In line with these complex neurobehavioral alterations, profound loss of central noradrenergic, serotonergic and thalamic neurons, and the development of synuclein aggregates have been described in chronically MPTP-treated monkeys and mice (Fornai et al., 2005; Masilamoni et al., 2011; Masilamoni et al., 2017; Masilamoni and Smith, 2018; Vivacqua et al., 2019). Another monkey study reported as much as 70% reduction in the number of catecholaminergic myenteric neurons (Chaumette et al., 2009); however, the effects of chronic systemic MPTP on *in vivo* gastrointestinal functions, inflammation, and bacterial composition have not been evaluated in non-human primates. With accumulating evidence highlighting the connection between the gut and the brain (van de Wouw et al., 2019), exploring the effects of nigrostriatal dopaminergic depletion on gastrointestinal symptoms will expand the relevance of these models in a multisystem disorder.

Increased levels of the pro-inflammatory cytokine tumor necrosis factor (TNF) have been described in postmortem brains and cerebrospinal fluid of PD patients (Eidson et al., 2017; Mogi et al., 1994) and in neurotoxin animal models of nigrostriatal degeneration (Barcia et al., 2005; Mogi et al., 1999), implicating its role in PD pathophysiology. Soluble TNF (sTNF) mediates pro-inflammatory responses by preferentially binding to the TNF receptor 1 (TNFR1) demonstrated by the increased susceptibility of bacterial infections in TNFR1 knockout mice (Rothe et al., 1993). Mice deficient in TNFR1 or both soluble and transmembrane TNF ligands are protected from MPTP-induced nigrostriatal degeneration (Ferger et al., 2004; Sriram et al., 2002), suggesting that TNF has a role in mediating neurotoxin-induced dopaminergic degeneration. Moreover, our group has previously shown that selective blocking of sTNF with XPro1595 was sufficient to protect against acute 6-hydroxydopamine neurotoxicity of dopaminergic neurons and neuroinflammation in rats (Barnum et al., 2014). Yet it is not clear whether nonmotor symptoms are tied to sTNF/TNFR1 signaling and whether XPro1595 would alter inflammation and microglial behavior in a progressive model of PD.

Thus, the primary objective of this study was to investigate the evolution of central and peripheral inflammatory responses elicited in a chronic low-dose MPTP monkey model of PD. Secondly, given the presence of nonmotor symptoms in PD and the need for appropriate pre-clinical animal models that replicate these features, we evaluated the gastrointestinal and cognitive behaviors in the slowly-progressing monkey model of PD. Finally, we evaluated the extent to which XPro1595 mitigated neuroinflammation and nonmotor symptoms elicited by MPTP. Importantly, during the course of the study, robust sex differences in inflammatory responses measured by PET imaging became apparent and later correlated very closely with an earlier onset of motor symptoms. Genetic studies have demonstrated that sex influences PD risk where the presence of a Y chromosome increases risk by 1.5 (Wooten et al., 2004). Additionally, sex-biased expression of genes related to neuron and immune function have been identified in nigral tissue of PD patients (Mariani et al., 2018). Yet the impact of sex on immune system responses and how the latter affect development of PD is not known. Therefore, this study provides evidence for the sex-specific influence on inflammatory responses in monkeys and underscores the importance of considering sex as a biological variable in preclinical animal models of PD.

## 2. Materials and Methods

### 2.1 Subjects and Ethics Statement

Five rhesus monkeys (*Macaca mulatta,* 5-7yrs of age, 3 females, 2 males) from the Yerkes National Primate Research Center (YNPRC) colony at Emory University were used in this study. All experimental procedures were approved by the Institutional Animal Care and Use Committee of Emory University and conducted in accordance with guidelines in the NIH Guide for Care and Use of Laboratory Animals (NIH Publications, 8^th^ edition, revised 2011). The monkeys were housed in temperature-controlled rooms and exposed to a 12-hour light cycle. Animals were fed twice daily with Lab Diet monkey chow supplemented with fruits and vegetables and water *ad libitum*.

### 2.2 Experimental Design

Following the collection of all baseline measures, monkeys were given weekly doses of intramuscular MPTP (0.2–0.8mg/kg, Sigma-Aldrich) starting at 0.2 mg/kg for 18 weeks and increasing to 0.8 mg/kg to maintain a parkinsonian state. All animals, regardless of sex, received the same dose by weight until stable parkinsonian motor symptoms appeared and were maintained, at which time MPTP administration was halted (Masilamoni et al., 2011). At 11 weeks, animals were randomly assigned to two groups (**Figure 1**), with consideration of sex, and peripheral subcutaneous treatment of XPro1595 (10mg/kg, n=2 female and n=1 male) or vehicle (n=1 female and n=1 male) began and continued every 3 days. Based on the clear differences in sensitivity to MPTP dosing between the 3 female and 2 male monkeys used in this study, once an animal in either sex group became mildly parkinsonian based on clinical rating, administration of MPTP was terminated to all animals of the same sex. These animals were followed for approximately 3-4 weeks to confirm the stability of the clinical status, and to conduct final endpoint procedures before euthanasia. Males received a total of 5.9 mg/kg of MPTP over 22 weeks (highest weekly dose was 0.5mg/kg) and were euthanized 26 weeks after the start of neurotoxin dosing, while females progressed more slowly than males and therefore received 15.2 mg/kg of MPTP over 37 weeks (highest weekly dose was 0.8mg/kg) and were euthanized at 40 weeks.

**Figure 1.**
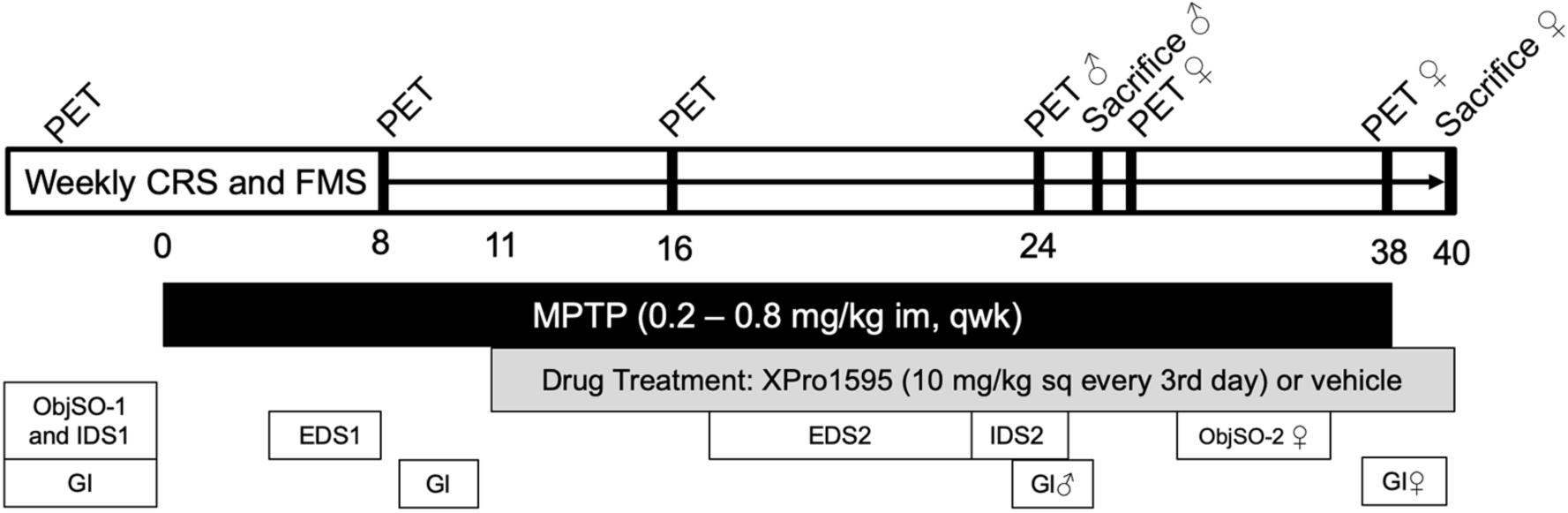
Experimental design and outcome measures. Animals performed a weekly fine motor skill (FMS) task and were assessed weekly for clinical parkinsonism status using a clinical rating scale (CRS). Weekly systemic peripheral MPTP injections (0.2 – 0.8 mg/kg, im) were administered to achieve and maintain mildly parkinsonian features by CRS. Drug treatment with either XPro1595 (10 mg/kg sq) or vehicle began 11 weeks after the start of MPTP injections and continued every 3 days until the end of the study. PET scans for [^18^F]FECNT and [^18^F]FEPPA were performed 1 week apart at the times denoted by “PET”. Bi-monthly blood and CSF samples were collected to analyze circulating markers of inflammation and levels of XPro1595. Cognitive behavior was tested throughout the study using an intradimensional (IDS) and extradimensional (EDS) shift task and an object self-order (ObjSO) task. Gastrointestinal measurements (GI) including functional analyses for permeability and specimen collection of colorectal biopsies and feces for microbiome analyses were conducted at baseline, ∼10 weeks after the start of MPTP treatment (prior to drug/vehicle treatment) and again at endpoint following either XPro1595 or vehicle treatment. Male and female monkeys were euthanized 26 and 40 weeks after the start of MPTP injections, respectively. Unless otherwise noted, procedures were done on both male and female monkeys.

### 2.3 Clinical and Motor Evaluations

Clinical status was assessed using a validated 27-point scoring system modified from the Unified Parkinson’s Disease Rating Scale (Masilamoni et al., 2011) by two investigators blinded to the treatment history of the groups. Each category was scored between 0-3 including: gross motor activity, posture, balance, bradykinesia (arm and leg), hypokinesia (arm and leg) and tremor (arm and leg). Assessments were begun after monkeys habituated to the observation cage for 15 minutes. An intra-class variability analysis between the two-experimenter scores demonstrated significant rater agreement with ICC of 0.977 (95% CI: 0.969-0.983). Gross motor behavior was assessed using an Actical accelerometer (Phillips Respironics, Bend OR) attached to a collar, while fine motor skills were assessed with a pickup test (FMS) (Masilamoni et al., 2011). Monkeys were placed into an observation cage once a week to perform the FMS in an apparatus with a clear plexiglass adhered to the front of the cage. The apparatus was fashioned with a center chamber where a treat was placed and portal holes for either the left or right limb to access the center chamber. Quantification was computed by the pickup test software and provided the time for the animal to pick up the treat in the center chamber and the total time to reach for the treat, retrieve it and retract the limb back into the cage.

### 2.4 Cognitive behavior

For all cognitive behavior, animals were acclimated to a Wisconsin General Test Apparatus (WGTA) contained in a sound-shielded room with a white noise generator to mask external sound. Animals were tested with an intradimensional/extradimensional (ID/ED) cognitive behavior task which requires monkeys to update their responses based on changing reinforcement contingencies (Weiss et al., 2019). Prior to MPTP, animals were tested on a visual compound discrimination training with a positive-rewarded stimulus and a negative-unrewarded stimulus. Following compound discrimination tests, animals were tasked with an intradimensional shift (IDS1; attention shift within same dimension) to measure associative learning. Five weeks after the start of chronic MPTP dosing and prior to administration of XPro1595 (or vehicle), animals were retrained to perform the compound discrimination followed by an extradimensional shift task (EDS1; attention shift to new dimension) to measure cognitive flexibility and a series of extradimensional reversals to measure behavioral inhibition. Following 6 weeks of XPro1595 (or vehicle) treatment and 17 weeks of escalating doses of MPTP, animals were retrained on a compound discrimination test and tasked with a new extradimensional shift (EDS2) and reversal. Finally, starting 10 weeks after the beginning of the XPro1595 (or vehicle) treatment, animals were retrained with a compound discrimination task and retested on an intradimensional shift (IDS2). In order to complete a stage and move onto the next, 10 consecutive correct trials had to be performed with up to 60 trials in a day, but a stage was failed if the criterion was not met in 500 trials. The number of trials and errors to reach criterion in each stage were used for analysis.

Animals were also tested with an object self-ordered task (ObjSO) to assess working memory and measure the monkey’s ability to monitor their own responses over a series of trials as a measure of working memory (Heuer and Bachevalier, 2011). Females were tested at baseline and starting at 30 weeks post-MPTP. A trio of unique objects were presented to the monkey in the WGTA. The animal displaced an object to retrieve a food reward. Next, the door was closed allowing the tester to rearrange the objects in a randomized fashion. The monkey was then given 3 trials with a 10-second inter-trial interval to displace an object with the goal of not repeating the placement of an object already selected. Primary errors were defined as the first incorrect choice on trial 2 or 3. Perseverative errors were defined as errors committed after a primary error on the same day. Criterion performance is 85% or better performance across 10 consecutive days of testing (i.e. 3 or less primary errors in 10 days), but task completion was deemed a failure if the criterion was not met in 50 sessions. The number of sessions to reach the criterion was recorded along with primary and perseverative errors during trials 2 and 3.

### 2.5 Gastrointestinal Assessments

A sugar permeability assay was conducted to determine intestinal barrier function; feces were collected for microbiome 16sRNA gene amplicon sequencing and colorectal biopsies obtained for inflammatory analysis at baseline, after 10 weeks of chronic MPTP administration and before drug treatment (MPTP) and prior to euthanasia after animals had been chronically challenged with MPTP and received either XPro1595 or vehicle treatment (MPTP + treatment). Additionally, lipopolysaccharide (LPS) and lipopolysaccharide binding protein (LBP) were evaluated from serum collected at baseline, 8 weeks after MPTP and prior to euthanasia as an indirect measure of circulating LPS due to gut leakiness.

#### Intestinal permeability assessment

GI permeability in humans has traditionally been measured by oral administration of a combination of sugars of different sizes: sucrose, lactulose, mannitol, and sucralose (Forsyth et al., 2011). The quantitation of the uptake and excretion of these substrates in urine samples allow for a relatively precise determination of the integrity of intestinal epithelial cell barrier function or disruption of barrier integrity resulting in passage/leak of these compounds into the blood stream and urine (McOmber et al., 2010; Shaikh et al., 2015). Animals were anesthetized with either Ketamine (5-10 mg/kg IM) or Telazol (3-5 mg/kg IM), intubated and then maintained under isoflurane (1-3%) anesthesia. A bladder catheter was placed to collect a baseline urine sample and to expel the urine remaining after baseline was established. After a baseline sample was collected, the animal was orally gavaged through a gastric tube with a bolus of up to ∼20 mL of a sugar solution (a combination of 40g sucrose, 7.5g lactulose, 2g mannitol and 2g sucralose per 70kg weight all dissolved in water). The animals remained anesthetized for ∼3 hours under the surveillance of the YNPRC veterinary staff. During this time, all urine was continuously collected via a bladder catheter. Three hours after administration of the sugar cocktail, the total urine volume from the bladder was measured and aliquots taken for gas chromatography analysis as previously described (Forsyth et al., 2011; Shaikh et al., 2015). Results are expressed as the amount of sugar excreted as a % of the oral dose.

#### Rectal biopsies and feces collection

Monkeys were anesthetized with either ketamine (5-10 mg/kg IM) or Telazol (3-5 mg/kg, IM) and placed in ventral or lateral recumbency. A lubricated anoscope was introduced in the rectum. Feces were removed and collected using sterile cotton-tipped applicators commonly used for stool collection for microbiome 16S rRNA gene sequencing and SCFA levels. Colorectal biopsies were obtained using a biopsy forceps under visualization by the Yerkes veterinarian staff. Both fecal and colonic biopsy samples were stored at −80°C until analyzed. Biopsies were later processed with homogenization buffer including a protease inhibitor (Roche) and 1% Triton X-100 and homogenized in a TissueLyser II (Qiagen), centrifuged at 20,000 rcf at 4°C for 10 minutes after which the supernatant was collected for multiplexed immunoassay analyses.

### 2.6 Radiosynthesis of [^18^F]FEPPA and [^18^F]FECNT

The preparation of [^18^F]FEPPA was similar to that previously described (Wilson et al., 2008). [^18^F]Fluoride was produced at YNPRC on a Siemens RDS 111 medical cyclotron, trapped and released, azeotropically dried, before undergoing a nucleophilic reaction with the tosylate precursor. The product was purified by reverse phase high-pressure liquid chromatography (HPLC) in buffered saline with 5-10% ethanol. The final product was passed through a 0.22-micron filter and formulated sterile, pyrogen free, to a pH of 5-8. Radiochemical purity was confirmed to be > 96% with specific activities greater than 1Ci/µmol.

The preparation of [^18^F]FECNT was conducted as previously described (Goodman et al., 2000; Masilamoni et al., 2010; Masilamoni et al., 2011). In the present study, we performed [^18^F]FECNT PET scans to determine the neuroprotective effects of XPro1595 on the nigrostriatal dopaminergic system during the course of escalating MPTP intoxication. *In vivo* PET imaging of dopamine markers is the most sensitive approach for longitudinal monitoring of the state of degeneration of the nigrostriatal dopaminergic system during the course of neuroprotective trials in PD (Pavese and Brooks, 2009). We have recently shown that the PET tracer [^18^F]FECNT, is a highly reliable ligand to estimate levels of striatal and nigral dopamine transporter in the MPTP-treated monkey model of PD (Masilamoni et al., 2010).

### 2.7 MRI scans

Animals were initially anesthetized with Telazol (3-5 mg/kg IM) and maintained using 1-2% isoflurane gas anesthesia for the duration of the imaging procedure. They were placed in a custom-made stereotaxic frame while in a supine position. A 3D T1-weighted MPRAGE (Mugler, 1999) of the brain was acquired with an Siemens Magnetom Trio 3-T (Siemens Medical Solutions USA, Malvern, PA) typically recommended for brain structure morphometric analyses (Lusebrink et al., 2013). The scan parameters were 0.5mm-thick images with a transverse plane isotropic pixel size of 0.5 mm (repetition time/echo time 2500/4.38 msec, inversion time of 900ms, flip angle of 10, acquisition field-of-view of 320 x 192 pixels, and a matrix of 320 x 320 pixels).

### 2.8 PET imaging

PET scans using [^18^F]FEPPA and [^18^F]FECNT were performed at baseline and at approximately 8 weeks (PET I), 16 (PET II), 24 (PET III) and 38 (PET IV) after the initiation of MPTP administration to monitor changes in microglial activation and dopamine transporter (DAT) binding, respectively. PET images were acquired in the YNPRC on a Siemens Focus 220 microPET scanner (Siemens, Concorde Microsystems, Knoxville, TN), which has an 8/26 cm axial/transaxial FOV and reconstructed image resolution is 1.7 mm in all directions. Animals were initially anesthetized with ketamine and Telazol then intubated and maintained on 0.8-1.5% isoflurane anesthesia for the duration of the imaging procedure. An acute venous catheter was placed in the saphenous vein for intravenous administration of the radioligand. The catheter access was maintained with a sterile saline drip throughout the study. Animals were placed supine in the micro-PET scanner, fitted with pulse oximetry equipment and a rectal thermistor for physiological monitoring during the procedure. Prior to injection of the PET radiopharmaceutical, a transmission scan was obtained with a Co 57-point source to correct for attenuation in the image reconstruction. Approximately 5 mCi bolus of radiopharmaceutical was injected intravenously over 5 min with an infusion pump set at a rate of 1.0 mL/min. The initial acquisition was a 28-frame dynamic sequence, starting with 30-sec scans and ending with 20-min scans (total duration, 115 min). PET data were reconstructed with a 3D Ordered subset Maximum Expectation algorithm including corrections for random, scatter, attenuation and dead-time.

### 2.9 PET data analysis

All images were decay-corrected to the time of injection. [^18^F]FECNT time-activity curves were generated for each monkey. PET images were co-registered to their structural MRI using IDL software (ITT Visual Information Solutions) and averaged to draw regions of interest (ROI) so that the volumes would be equal and comparable over all treatment time points. ROIs were manually drawn as defined by the Paxinos atlas including the following striatal regions: putamen/associative (PA), putamen/motor (PM), putamen/limbic (PL), caudate nucleus/associative (CA), nucleus accumbens/limbic (AC) and the substantia nigra (SN) (Paxinos et al., 2000). The regions of interest were then superimposed onto the individual animal PET images to obtain time-activity curves. Because of the minimal expression of dopamine transporter in the cerebellum, we used it as the reference region, and expressed the FECNT uptake value as the non-displaceable binding potential, as described in our previous studies (Masilamoni et al., 2010; Votaw et al., 2002).

For quantification of [^18^F]FEPPA, regions of interest were manually drawn and defined on the MRI within structural boundaries defined by the Paxinos atlas (Paxinos et al., 2000), including the cerebellum, frontal cortex, parietal cortex, occipital cortex, temporal cortex, putamen, substantia nigra, and lateral geniculate nucleus. The manually-drawn ROIs were superimposed onto the individual animal [^18^F]FEPPA PET images to obtain time-activity curves using IDL software (ITT Visual Information Solutions). [^18^F]FEPPA uptake was expressed as Standardized Uptake Values (SUV), accounting for body mass and injected activity of the tracer (Zhang et al., 2012). SUV allows precise assessment of activity changes in response to drug treatments (Masilamoni et al., 2016; Zhang et al., 2012). No reference brain region was used to assess background activity because systemic MPTP administration is likely to affect all regions of the brain, albeit differentially. [^18^F]FEPPA uptake was validated using a LPS-treated monkey model (**Figure S1**). The total binding was determined by calculating the mean SUV for [^18^F]FEPPA data from 85 – 115 minutes post-injection.

### 2.10 Blood collection

Animals were anesthetized with ketamine (10 mg/kg, IM) to collect blood and CSF monthly. Blood was collected in EDTA tubes from either femoral or saphenous veins. Plasma was extracted after a 15-minute centrifugation at 3000rpm at 4°C. CSF was obtained by lumbar puncture using sterile technique. All samples were stored at −80°C until assayed. Additional blood was collected at baseline, 8 weeks after MPTP and at endpoint to evaluate LPS and LPS binding protein (LBP) in a serum separator tube, allowed to remain at room temperature for at least 30 minutes to clot, and serum extracted after a 10-minute centrifugation at 1000g. Animals’ vital signs were monitored by YNPRC veterinary staff until animals fully recovered.

### 2.11 Necropsy and Tissue Preparation

Male and female animals were anesthetized with ketamine (10 mg/kg IM) and sodium pentobarbital (100 mg/kg IV) 26 or 40 weeks after the start of MPTP toxin dosing, respectively. The descending aorta was clamped and animals were trans-aortically perfused with cold oxygenated Ringer’s solution followed by 4% paraformaldehyde (PFA) and 0.1% glutaraldehyde in phosphate buffer (0.1M, pH 7.4) (Masilamoni et al., 2011). Prior to perfusion with PFA, a burr hole was drilled into the skull and a cortical biopsy taken using a skin biopsy tool for measurements of XPro1595 brain levels. The brain was removed and placed in 4% PFA for 48 hours and then cryoprotected in 30% sucrose. Sections of each segment of the small and large intestine were post-fixated in 4% PFA for 48 hours and then incubated in 70% isopropyl alcohol. Gut sections were processed and blocked in paraffin, cut on a rotary microtome at 5-micron thick sections and mounted on positive-charged glass microscope slides. The right hemisphere of the brain was cut in the coronal plane on a freezing microtome into 50-micron thick sections. All tissue was stored in cryoprotectant at −20°C before being processed for immunohistochemical or immunofluorescence analyses.

### 2.12 Multiplexed immunoassays

Plasma, CSF and gut biopsies were analyzed for IFN-γ, IL-10, IL-1β, IL-2, IL-6, and IL-8 using a non-human primate V-PLEX pro-inflammatory panel (K15056D-1 MesoScaleDiscovery, Rockville, MD) multiplexed immunoassay with 1:2 dilutions for plasma, and biopsies assayed at 2μg/μL. Biofluids were also measured for neutrophil gelatinase-associated lipocalin (NGAL), C-C motif chemokine ligand 2 (CCL2; also known as monocyte chemoattractant protein-1, MCP1) and TIMP-1 using a 3-PLEX cynomologous (catalog #K1519D-1, MSD) with 1:20 dilution for plasma, and for C-reactive protein (CRP) using a single analyte immunoassay (catalog #K151STD) with 1:1000 dilution for plasma. CSF was analyzed using 1:1 dilution. XPro1595 was measured using an anti-human TNF ultrasensitive immunoassay (catalog#K151QWD, MSD) for brain biopsies at 2μg/μL, plasma diluted 1:300 and non-diluted CSF. All samples were measured in duplicate and immunoassays were performed as per the manufacturer’s instructions by the Emory Multiplexed Immunoassay Core.

### 2.13 Immunohistochemistry and Immunofluorescence

Brightfield immunological staining of coronal brain sections was performed as described previously (Masilamoni et al., 2011; Mathai et al., 2015) using antibodies against tyrosine hydroxylase (RRID:AB_572268; 1:10,000, Immunostar, Hudson WI), and macrosialin/CD68 clone KP1 (RRID:AB_10987212; 1:400, Thermofisher, Waltham, MA). Tissue sections were further incubated in biotinylated secondary antibodies raised against the appropriate species, followed by avidin-biotin complex and developed in solution containing 0.025% 3,3’-diaminobenzidine tetrahydrochloride (Sigma-Aldrich), 10 mM imidazole, and 0.005% hydrogen peroxide in Tris buffer. CD68 staining was enhanced by the addition of nickel sulfate. Slides containing gut tissue were deparaffinized in xylene and rehydrated before performing antigen retrieval as previously published (Joers et al., 2014; Shultz et al., 2016). Gut tissue was similarly stained for CD68 without nickel enhancement.

For immunofluorescent staining, free-floating tissue was blocked in 5% serum, 1% bovine serum albumin and 0.3% Triton-X-100 for 1 hour followed by primary antibody incubation overnight at room temperature with an antibody specific for microglia markers IBA-1 (RRID:AB_839504; Wako, Richmond VA) and B-lymphocyte LN-3 (HLA-DR; catalog 0893031, MP Biomedical, Santa Ana, CA). Sections were next incubated in secondary antibodies conjugated to FITC and rhodamine red (Jackson ImmunoResearch Inc, West Grove, PA) diluted to 1% for 1 hour at room temperature. Non-specific binding was diminished with a 30-minute incubation in cupric sulfate solution at pH of 5.0. Tissue was mounted onto coated glass slides and coverslipped with Vectashield mounting media (Vector laboratories, Burlingame, CA).

### 2.14 Stereological estimation of TH+ and Nissl cells

The unbiased stereological estimation of the number of dopamine neurons in the ventral substantia nigra pars compacta (SNv), dorsal substantia nigra pars compacta (SNd), ventral tegmental area (VTA), locus coeruleus (LC) was achieved using the optical fractionator probe (StereoInvestigator, MicroBrightField, Inc.), a stereological approach that combines the optical dissector with a fractionator sampling scheme. This sampling technique is not affected by tissue volume changes and does not require reference volume determinations. The random systematic sampling of counting areas was done using the Leica DMR microscope. TH immunoreactivity and Nissl staining was used to sample cell number; these markers provide information as to whether reductions in the numbers of TH + cell bodies are due to neuronal destruction or to a loss or downregulation of TH phenotype in the surviving neurons. To quantify midbrain TH+ and Nissl+ neuron number, slides were scanned at 20x using a ScanScope CS scanning system (Aperio Technologies, Vista, CA). Digital representations of the slides were saved and viewed using ImageScope software (Aperio Technologies). Low-power micrographs (1.8x) of TH, Nissl and calbindin-immunostained ventral midbrain sections were used to manually delineate the borders of the SNv, SNd, and ventral tegmental area, based on the presence or absence of calbindin-positive neurons (Masilamoni et al., 2010). Then, the borders of the different ventral midbrain regions were manually delineated on TH-immunolabelled and Nissl-stained slides adjacent to those immunostained for calbindin. Counts of TH+/Nissl+ cells were generated using a 100x oil-immersion objective. To perform unbiased stereology, counting frames (65 × 65 µm) were randomly placed by the stereology software within the chosen ROI. The software also controlled the position of the x–y stage of the microscope with a grid size of 300 x 300-µm using a dissector height of 27-µm and a 2-µm guard zone, so that the entire brain region could be scanned; 11 SN sections and 7 locus sections were used for stereological analyses. The software algorithm calculated the estimated total number of cells in each region of interest per hemisphere. The analysis was performed by one examiner blind with regard to the experimental condition.

### 2.15 Analysis of microglia number and phenotype

Analysis of IBA1 and HLA-DR was conducted in the area of the SN. A ROI was placed around the DAT-expressing area and imaged using automatic tiling on an Olympus FV1000 confocal microscope at 20x (N.A 0.75, z-step: 1.00 μm). Within the SN ROI, 5 fluorescent confocal stacks were collected at random for each piece of nigral tissue stained for DAT, IBA1 and LN3 (n=2 per animal). High-magnification image stacks were imported into Imaris (version 7.2.2) and IBA1+ cells were identified using the “spot” function and assigned a designated number. Then, using a random number generator, a microglia cell was randomly selected from each image stack and a surface reconstruction performed of the IBA1 and HLA-DR based on fluorescence threshold. For quantification, the surface area and volumes of individual microglia were summed across all images and the percent coverage of HLA-DR per cell was calculated.

### 2.16 Analysis of CD68-immunoreactivity in gastrointestinal samples

Two sections from the Ileum and descending colon were evaluated for CD68-immunoreactivity. Microphotographs were collected from two sites with intact villi avoiding those with large lacteal structures in the ileum and from colonic crypts in both the longitudinal and cross-section orientation. Two villi and longitudinal crypts per image were outlined, while a 100-mm^2^ ROI was positioned over cross-section crypts for analysis. Images were analyzed using ImageJ software (version 1.52e) for optical density and area above threshold and averaged within animal.

### 2.17 Microbiota Profiling and Bioinformatics Analysis

Total DNA was extracted from the monkey fecal samples utilizing the FastDNA bead-beating Spin Kit for Soil (MP Biomedicals, Solon, OH, USA), and DNA concentrations verified with fluorometric quantitation (Qubit, Life Technologies, Grand Island, NY, USA). Sequencing libraries were generated using a two-stage “targeted amplicon sequencing (TAS)” protocol, as described previously (Naqib et al., 2018). Briefly, genomic DNA was PCR amplified with primers CS1_515F and CS2_806R (modified from the primer set employed by the Earth Microbiome Project (EMP; GTGYCAGCMGCCGCGGTAA and GGACTACHVGGGTWTCTAAT) targeting the V4 region of microbial small subunit ribosomal RNA genes. Subsequently PCR products were amplified using Access Array Barcodes for Illumina primers (Fluidigm, South San Francisco, CA) to incorporate Illumina sequencing adapters and sample-specific barcodes. Sequencing was performed using an Illumina MiSeq (Illumina, San Diego, CA, USA), with a V3 kit at the Sequencing Core at the University of Illinois at Chicago (UICSQC). Raw sequence data (FASTQ files) were deposited in the National Center for Biotechnology Information (NCBI) Sequence Read Archive (SRA), under the BioProject identifier PRJNA597903.

Raw FASTQ files for each sample were merged using the software package PEAR (Paired-end-read merger) (v0.9.8)(Schmieder and Edwards, 2011; Zhang et al., 2014). Merged reads were quality trimmed, and primer sequences removed. Sequences shorter than 225 base pairs were discarded. Sequences were screened for chimeras (usearch8.1 algorithm), (Edgar, 2010) and putative chimeric sequences were removed from the dataset (QIIME v1.8) (Caporaso et al., 2010). Each sample was rarefied (34,000 sequences/sample) and data were pooled, renamed, and clustered into operational taxonomic units (OTU) at 97% similarity (usearch8.1 algorithm). Representative sequences from each OTU were extracted and classified using the uclust consensus taxonomy assigner (Greengenes 13_8 reference database). A biological observation matrix (BIOM)(McDonald et al., 2012) was generated at each taxonomic level from phylum to species (“make OTU table” algorithm) and analyzed and visualized using the software packages Primer7 and the *R* programming environment (Clarke, 1993; R Development Core Team, 2013). Following sequencing, computational bioinformatics analysis was used to describe the bacterial diversity, community structure, and composition of the microbiota, as described previously (Bishehsari et al., 2018; Mahdavinia et al., 2018; Xiao et al., 2019). Dominant phylum, family, and genus bacteria were determined using a relative abundance threshold >2%, within the sex of monkeys. Both highly abundant (>2%) and low abundant (<2%) individual taxa were examined.

### 2.18. Short chain fatty acids levels: a marker of microbiota function

For SCFA measurements, stool was weighed, mixed with 3 parts carbonate-phosphate buffer and homogenized, and it was then centrifuged at 13000 rpm for 5 min. A mixture of 5 % phosphoric acid containing 50mM 4-methylvaleric acid and 1.56 mg/mL copper sulfate in water was prepared as internal standard and was added to the supernatant. 4ul of sample was injected onto a capillary column in an Agilent 6890 GC with a flame ionization detector as described previously (Bishehsari et al., 2019; Y.E. et al., 2017). The GC column used was a Nukol (Supelco Bellefonte; PA) 30m length, inner diameter 0.25-mm ID, 0.25-µm bonded phase. The GC was run under the following conditions: injector temperature 240°C; detector temperature 230°C.The initial oven temperature 100°C was held for 10 min and then increased at 8°C to 192°C for total run time of 22 minutes. The carrier gas helium was maintained at 1ml/min flow rate.

### 2.19 Statistical Analysis

Data was statistically analyzed using Graphpad software (version 6f). A two-way ANOVA was conducted to evaluate inflammatory markers across time against sex and a Sidak’s multiple comparisons performed. All correlations were performed using a Pearson’s correlation coefficient. Microglia histology was not statistically analyzed due to the limitations when separating sexes within treatment groups. Slopes of a linear regression analysis were compared between treatments for inflammatory markers to determine the divergence between groups. For nonmotor evaluations prior to drug treatment, a two-way ANOVA was first performed to evaluate the effect of sex on the outcome measure. If sex did not statistically influence the results, animals were grouped together, and a Student’s t-test was used to analyze the difference between baseline and early MPTP time points. Otherwise, the two-way ANOVA results were reported to demonstrate sex-specific effects. The % change from baseline was used to evaluate differences between MPTP and MPTP + drug treatment for gastrointestinal measurements including alpha diversity and SCFA concentrations.

## 3. Results

### MPTP-induced microglia activation occurred earlier in males resulting in sex-specific patterns of TSPO expression

Neuroimaging was employed to monitor *in vivo* microglial activation and dopaminergic neuron health. TSPO expression was monitored as a proxy for inflammatory activity and related to DAT expression as a marker of dopaminergic function on PET scans with [^18^F]FEPPA and [^18^F]FECNT ligands respectively, at baseline and after weekly MPTP injections (3 post-MPTP scans for males, and 4 for females, **Figure 2**). Consistent with our previous studies (Masilamoni et al., 2010; Masilamoni et al., 2011), [^18^F]FECNT binding in the putamen motor and SN acutely increased after the start of MPTP and plummeted by endpoint scans in both males and females, a pattern that was not affected by XPro1595 treatment (**Figure S2**). Both male monkeys demonstrated a robust increase in [^18^F]FEPPA binding (91% above baseline) in the putamen motor after 8 weeks of MPTP dosing prior to the start of XPro1595/vehicle treatment (PET I). After drug treatment, both male monkeys demonstrated a global progressive decline of [^18^F]FEPPA binding in the putamen motor throughout the remainder of the study (**Figure 2E-F**). This pattern of [^18^F]FEPPA binding was markedly different from that seen in the three female monkeys (**Figure 2G-H**). Instead, the average group [^18^F]FEPPA binding in the putamen motor remained close to baseline levels after 8 weeks of MPTP initiation (PET I), with one female displaying a marginal increase in TSPO levels (37% above baseline). In contrast, [^18^F]FEPPA binding in female monkeys dropped to or below baseline values after MPTP administration began and gradually increased by the endpoint scan (PET IV). There was no distinction of [^18^F]FEPPA binding across time between the XPro1595- and vehicle-treated animals when data from males and females were analyzed separately. Similar sex-specific patters of [^18^F]FEPPA binding were found in the SN (**Figure 2F,H**).

**Figure 2.**
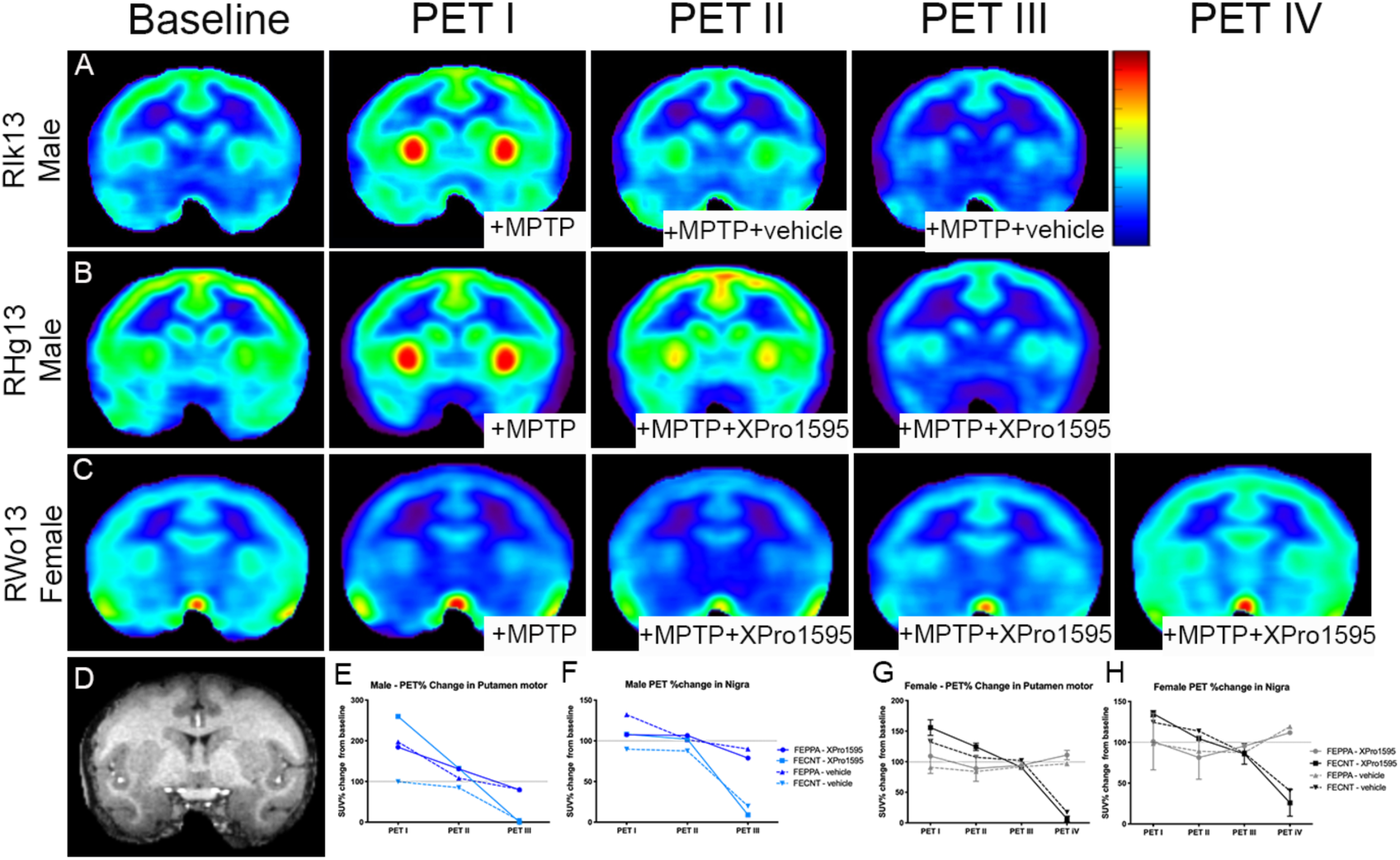
[^18^F]FEPPA PET reveal sex differences in response to chronic systemic MPTP injections. Representative coronal images of [^18^F]-FEPPA uptake in a post-commissural striatal slice of chronically MPTP-treated male and female rhesus monkeys treated systemically with vehicle (A) or XPro1595 (10mg/kg, B,C) at baseline, 8 (PET I), 16 (PET II), 24/27 (PET III) and 39 weeks (PET IV) after the start of MPTP administration. Coronal MRI image of the post-commisural striatum that matches the plane of PET images (D). [^18^F]FEPPA quantification as it relates to [^18^F]FECNT binding in the putamen and substantia nigra of males (E,F) and females (G,H) shows distinct patterns of [^18^F]FEPPA binding between sexes with no discernable effect of XPro1595 treatment. In males, [^18^F]FEPPA binding increased immediately after MPTP dosing (PET I) and globally declined overtime, following a trajectory similar to the [^18^F]FECNT % change. In females, [^18^F]FEPPA binding remained close to baseline levels through most scans, except for a slight increase in PET IV, at time point corresponding to severe decrease [^18^F]FECNT in the putamen motor.

In addition, the relationships between the pattern of [^18^F]FECNT and [^18^F]FEPPA binding differed between males and females. While the two ligand bindings in both the striatum and SN were closely related in the male monkeys (**Figure 2E-F**), in female monkeys the PET IV revealed opposite relationships between the slight increase in [^18^F]FEPPA and the robust decrease in [^18^F]FECNT binding (**Figure 2G-H**).

Consistent with an earlier and more robust microglia response, male monkeys exhibited a significantly higher level of endogenous TNF in the CSF than females after 4 weeks of MPTP dosing, despite having received the same amount of MPTP based on weight (**Figure 5C**). Due to the unexpected and robust sex-specific effect of MPTP on microglial responses and onset of parkinsonism, we were compelled to stratify the groups by sex for analysis of drug effects on MPTP treatment.

### Postmortem histological analyses of microglial activation corroborate the PET findings

Postmortem, we used Imaris reconstruction to quantify immunofluorescence staining of IBA1 and HLA-DR to assess microglial phenotype and related these observations to the last [^18^F]FEPPA PET scan results from the SN (**Figure 3**). The averaged surface area of reconstructed nigral IBA1+ microglia was significantly correlated with nigral [^18^F]FEPPA SUV (R^2^=0.837, p=0.029, **Figure 3K**), but not with the antigen presentation marker HLA-DR (data not shown). The nigral IBA1 immunostaining surface area was greater in female monkeys than in historical sex-matched controls and males used in this study (**Figure 3H**). Additionally, the male monkey treated with XPro1595 demonstrated an appreciable reduction in IBA1+ surface area and volume compared to its vehicle counterpart (**Figure 3H-I**). Of note, the surface area ratio of HLA-DR to IBA1 in the male that received XPro1595 treatment was different from what was found in females that underwent the same drug treatment (**Figure 3J**).

**Figure 3.**
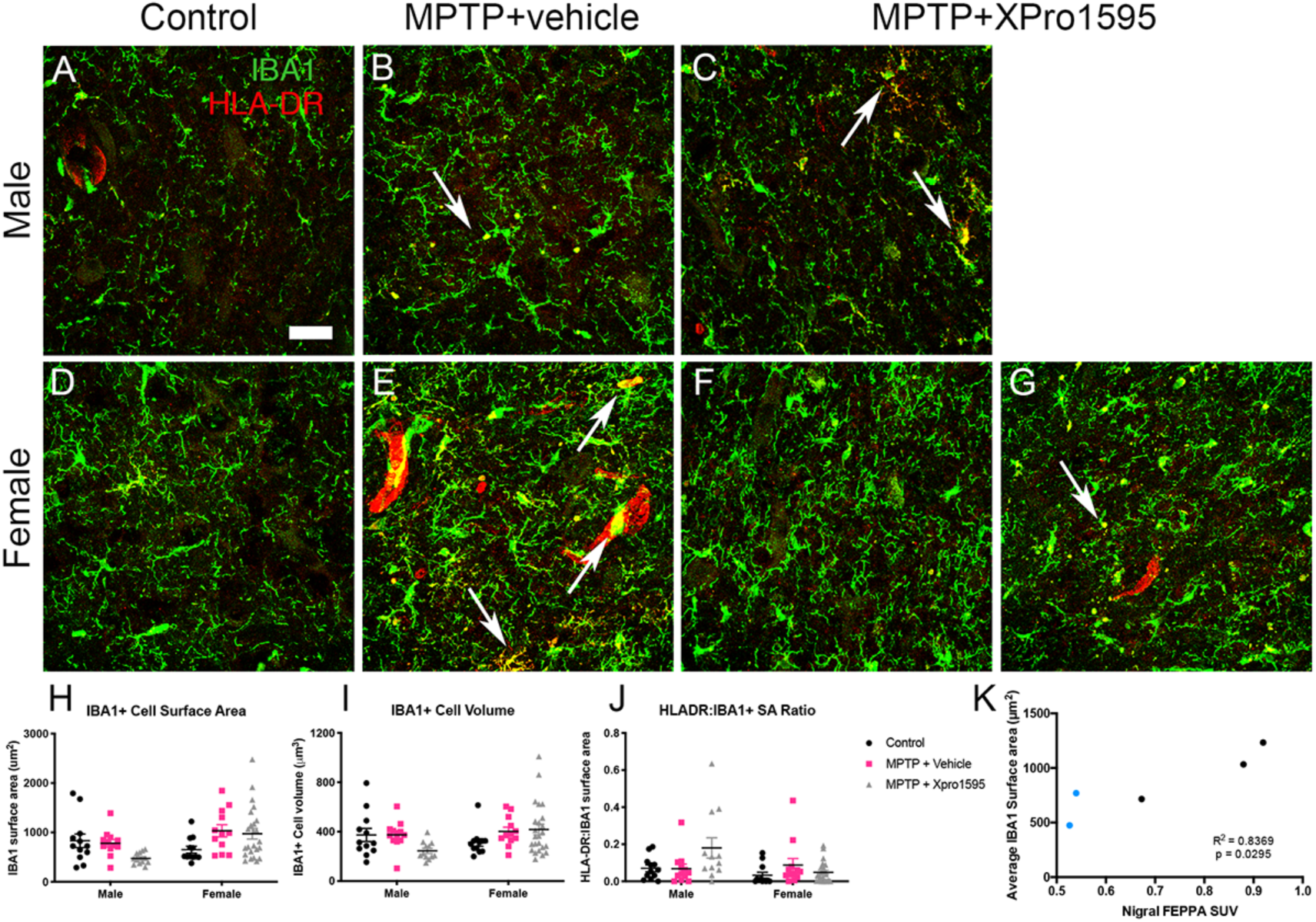
Histological analysis of IBA1 corroborates neuroimaging FEPPA analysis. Confocal images of IBA1 (green) and HLA-DR (red) in male and female controls (A,D), MPTP and vehicle-treated (B, E) and MPTP/XPro1595-treated male and female monkeys (C,F-G). Evaluation of reconstructed nigral IBA1+ microglia (n=12/animal from a rostral and caudal nigral section) highlights the differences in IBA1 surface area (H), cell volume (I) and HLA-DR/IBA1 surface area ratio (J) between male and female monkeys treated with XPro1595. Significant correlation between IBA1 surface area and nigral [^18^F]FEPPA binding (R^2^=0.837, p=0.029). Males are represented by blue points (K).

We further investigated the microglial phenotype using immunohistochemical staining of the phagocytic and lysosomal marker CD68 also known as macrosialin (**Figure 4**). Quantification of CD68+ particle counts highlighted an effect of MPTP compared to historical controls and an effect of drug treatment that was strongest in females. As demonstrated by the noticeable increased number of CD68+ particles in XPro1595-treated females compared to the vehicle-treated female monkey, which was not seen in the cohort of male monkeys (**Figure 4C**).

**Figure 4.**
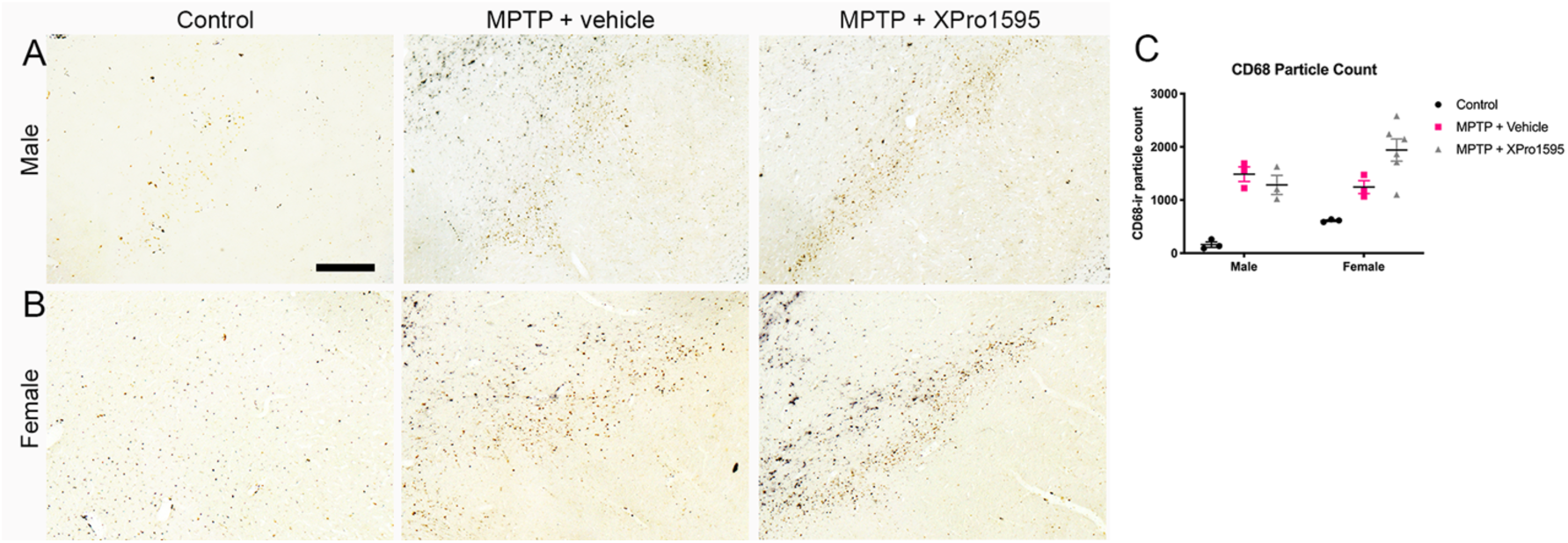
CD68-ir increases in chronic MPTP-treated monkeys. Representative CD68-stained nigral sections with nickel (black staining) from a male (A) and female (B) control, MPTP/vehicle- and MPTP/XPro1595-treated monkeys. Increase of CD68-ir particle counts from 3 regions in a rostral and caudal section of the nigra in all MPTP-treated monkeys compared to historical controls. XPro1595 treatment resulted in increasing CD68+ counts in females compared to vehicle-treated animals and little effect in males of different treatments (C). Scale bar 500 µm.

### Sex differences in MPTP-induced parkinsonism and immune responses

Following weekly administration of escalating doses of MPTP, two patterns of sensitivity to the neurotoxin emerged dependent on sex and independent of drug treatment. Male monkeys developed earlier parkinsonism as measured by motor deficits and were sacrificed at 26 weeks, while females developed parkinsonian symptoms much later and were sacrificed at 40 weeks (**Figure 5A**). A two-way ANOVA analysis of the grouped row means of CRS over time demonstrated no significant difference between XPro1595 treatment and vehicle groups at any timepoint (F(1,94)=0.546, p=0.462). The escalating delivery of weekly MPTP led to a significant change of clinical status over time independent of treatment (F(39, 94) = 3.927, p<0.0001). CRS impairment was echoed in the fine motor assessments using the FMS task, such that animals with the highest clinical score exhibited slower times to retrieve a treat (**Figure S3**). Similar to the pattern in individual clinical status, plasma IL-6 increased at 12 weeks and continued to increase with escalating MPTP doses independent of treatment and sex (**Figure 5B**). Given the distinct sex differences in MPTP sensitivity and uneven timelines, the relationship between inflammatory mediators and time was further evaluated by sex (**Figure 5D-I**). A significant interaction was found between the effects of sex and time for NGAL CSF (F(1,3)=26.2, p=0.014), and plasma NGAL (F(1,3)=26.2, p=0.014) and CRP (F(7,21)=10.21, p<0.0001) such that males presented with increased inflammatory levels compared to the females. Levels of CSF IL-6 (F(7,21)=14.52, p<0.0001), IFNγ (F(7,23)=4.71, p=0.0021), and plasma IL-8 (F(7,21)=5.276, p=0.0014) significantly increased over time regardless of sex.

**Figure 5.**
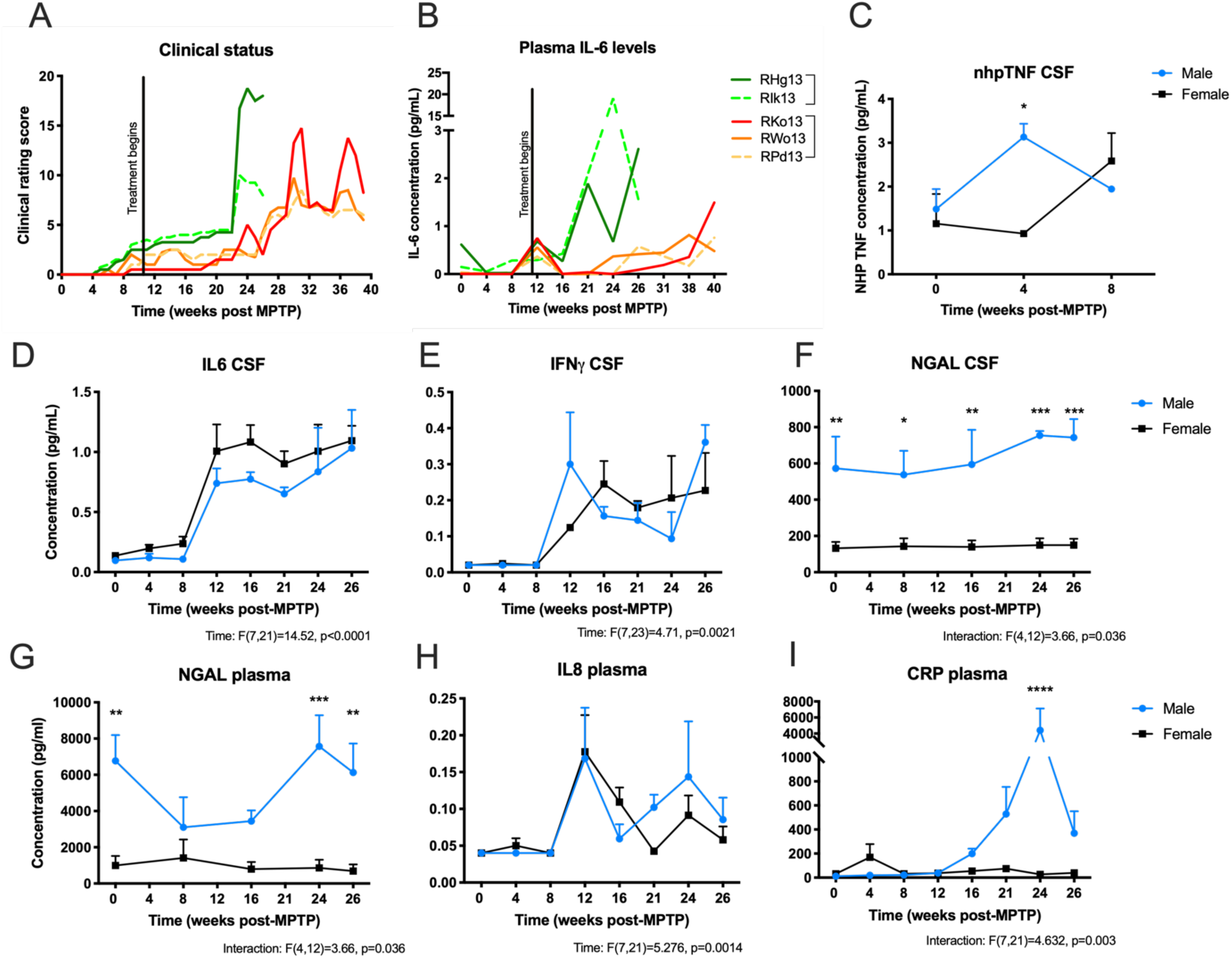
Chronic MPTP induces sexually dimorphic parkinsonism and immune responses in monkeys. Animals were evaluated *in vivo* with a validated modified Unified Parkinson’s Disease Rating Scale or clinical rating score (CRS) following weekly MPTP administration. Individual assessment of CRS demonstrates increasing impairment and a sex difference in sensitivity to MPTP that was not affected by XPro1595 treatment (A). Plasma IL-6 levels show the same pattern as CRS revealing increased levels in late disease states (B) that was also correlated to CRS (R^2^(48)=0.1498, p=0.0066). Dashed lines in A and B represent vehicle-treated animals and solid lines represent XPro1595-treated animals. Levels of CSF nhpTNF were increased in males but not in females at 4 weeks after the start of MPTP treatment (C). Cytokine levels stratified by sex across time reveal a significant MPTP or sex-specific effects on CSF IL-6 (D), IFNγ (E), and NGAL (F) and plasma NGAL (G), IL-8 (H) and CRP (I).

### Delayed administration of XPro1595 mitigated pro-inflammatory markers in biofluids but did not protect against nigral degeneration

Due to the nature of the model used herein where MPTP doses were slowly escalated over time, chronically administered MPTP led to an expected significant reduction in [^18^F]FECNT at the final PET timepoint, TH+ nigral and locus coeruleus cell death as determined by stereological counting, and reduced putamen TH immunoreactivity as measured by optical density as compared to historical controls (**Figure S4**). Because no significant sex differences in TH deficits were detected between sexes in the SN (F(1,9)=0.723, p=0.417) or the lower brainstem noradrenergic cell groups (F(1,9)=0.299, p=0.867), data from males and females were grouped together for analysis across treatment. Similar patterns of TH immunoreactivity loss were found in both MPTP-treated groups, apart from a significantly lower number of TH+ neurons in the SNCv of the XPro1595-versus vehicle-treated animals (p<0.05).

Taking note of the inflammation measured throughout the study, a positive relationship was found between various cytokine levels and CRS, suggesting that increased levels of inflammation accompany motor impairment (**Figure 6**). Specifically, vehicle-treated animals CRS correlated with CSF immune mediators CRP (R^2^(11)=0.339, p=0.037) and NGAL (R^2^(7)=0.719, p=0.0039), and plasma immune mediators IL-6 (R^2^(11)=0.475, p=0.009), CRP (R^2^(11)=0.39, p=0.023), and CCL2 (R^2^(8)=0.503, p=0.022), whereas XPro1595-treated monkeys demonstrated a significant association of clinical rating score with CSF levels of NGAL (R^2^(13)=0.267, p=0.0485) and plasma levels of IL-6 (R^2^(19)=0.228, p=0.029), CRP (R^2^(19)=0.242, p=0.023) and CCL2 (R^2^(13)=0.429, p=0.008). Notably, the slopes of best fit lines between the vehicle- and XPro1595-treated animals were significantly different when comparing the relationship between the CRS and CSF IL-6 (F(1,29)=4.762, p=0.0373) and plasma IL-6 (F(1,30)=20.06, p=0.0001) and CCL2 (F(1,21)=16.55, p=0.0006). Importantly, significant differences in slope between treatment groups indicate that XPro1595 treatment dampened inflammation. Systemic dosing of XPro1595 achieved plasma levels of 10-117μg/mL, CSF levels of 3-83 ng/mL, and cortex levels of 1-38 ng/mL in animals that received the drug, while vehicle animals were below detectable levels or below the lowest levels found in XPro1595-treated animals (**Figure S5**), consistent with previous published studies in rodents (Barnum et al., 2014; MacPherson et al., 2017). Additionally, no sex differences in systemic or central XPro1595 levels were observed.

**Figure 6.**
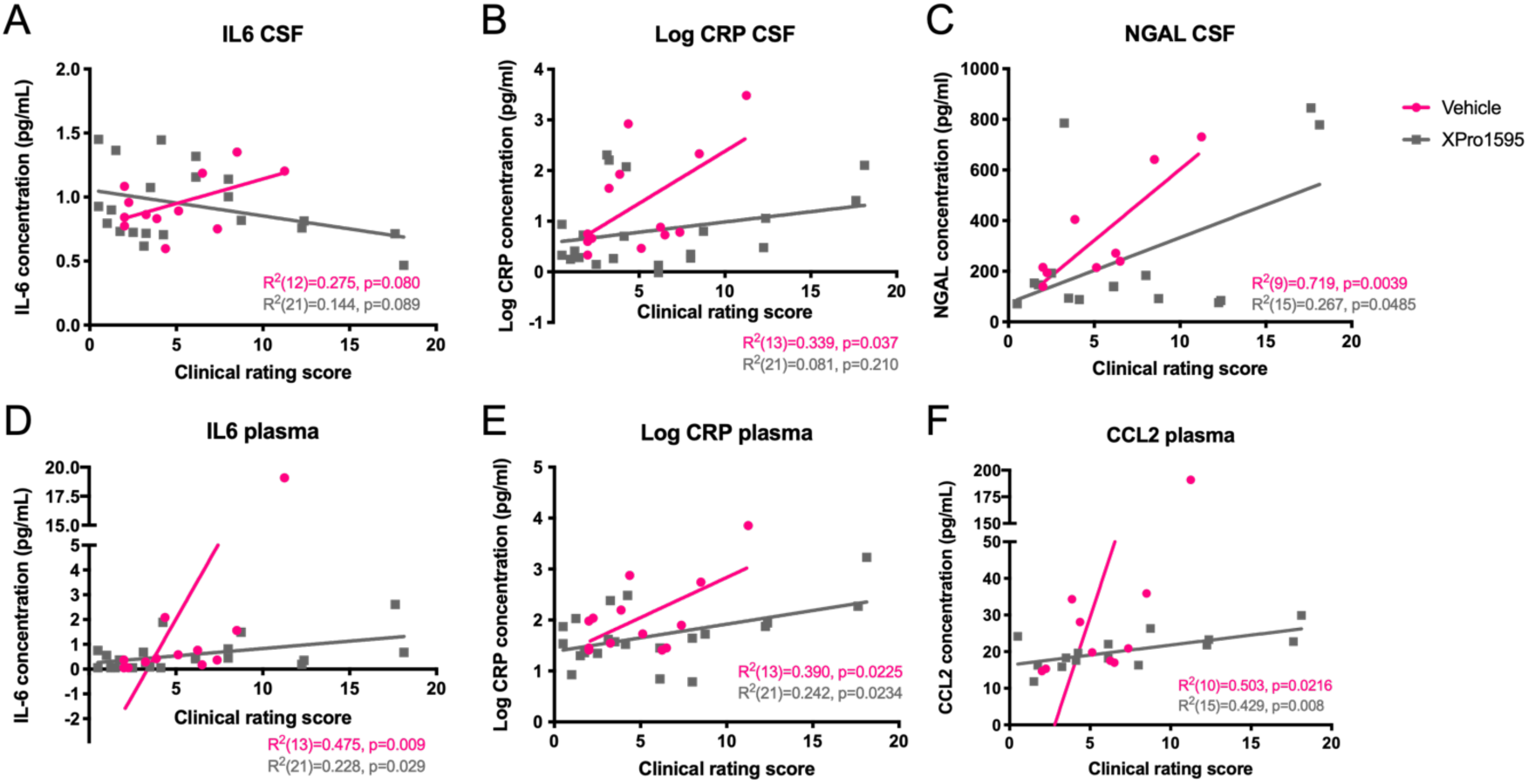
XPro1595 shifts MPTP-induced central and peripheral inflammation. When the relationship between clinical status and inflammation markers are stratified by treatment, there is a distinct divergence in the levels of cytokines in biofluids between XPro1595- and vehicle-treated MPTP intoxicated monkeys. In vehicle-treated animals, significant correlations were found between CRS and CSF IL-6 (A), Log CRP (B), NGAL (C) and plasma IL-6 (D), Log CRP (E), CCL2 (F), whereas XPro1595-treated animals display such correlations only for plasma CRP and CCL2. The slope of the best fit line is significantly shifted downward in XPro1595-treated animals for CRS correlation with CSF IL6 and in plasma IL6 and CCL2. IL-6 CSF: <u>F(1,29)=4.76, p=0.037</u>; CRP CSF: F(1,30)=3.72, p=0.063; NGAL CSF: F(1,20)=1.35, p=0.260. IL-6 plasma: <u>F(1,30)=20.06, p=0.0001</u>; CRP plasma: F(1,30)=3.08, p=0.089; CCL2 plasma: <u>F(1,21)=16.55, p=0.0006.</u>

### Chronic MPTP administration impaired cognitive flexibility and increased perseverative errors but maintained intact associative learning

In addition to motor behavior deficits, chronic low doses of MPTP independent of drug treatment resulted in impaired cognitive flexibility (EDS stages) measured by an increased number of errors to reach criterion (**Figure 7A-C**; F(1,3)=11.38, p=0.043). Animals experienced difficulty performing EDS reversals in both early (EDS1; 5 weeks after MPTP) and late (EDS2; 17 weeks after MPTP) MPTP stages. In the early MPTP dosing stage, only 2 females (one from each treatment group) successfully reached criterion for 3 reversal stages, while in the late MPTP dosing stage, the vehicle-treated female was the only one that successfully passed the EDS2 and one reversal (data not shown). However, monkeys displayed intact (4 of 5 animals) stimulus-response association learning after at least 22 weeks of MPTP dosing (IDS stages; **Figure 7D-F**). Due to the shortened timeline to parkinsonism for the males, only female monkeys were evaluated for the ObjSO task post-MPTP administration. Following MPTP and treatment, all three females demonstrated impaired performance on the ObjSO task with increased number of trials to reach criterion compared to baseline (t(2)=5.55, p=0.031) and increased primary (t(2)=4.76, p=0.041) and perseverative errors in the third trial post-MPTP compared to baseline (t(2)=9.938, p=0.01) with no difference between the treatment groups (**Figure 7G-I**). Deficits in EDS task correlated with central levels of CSF in both males and females (R^2^=0.534, p=0.0164; **Figure 7J**). Additionally, both primary and perseverative errors positively correlated with CSF levels of IL-6 (Trial 2 primary R^2^=0.899, p=0.0039; Trial 3 primary R^2^=0.8181, p=0.0132; Trial 3 perseverative R^2^=0.845, p=0.0095; **Figure 7K**), indicating a relationship between central inflammation and cognitive performance.

**Figure 7.**
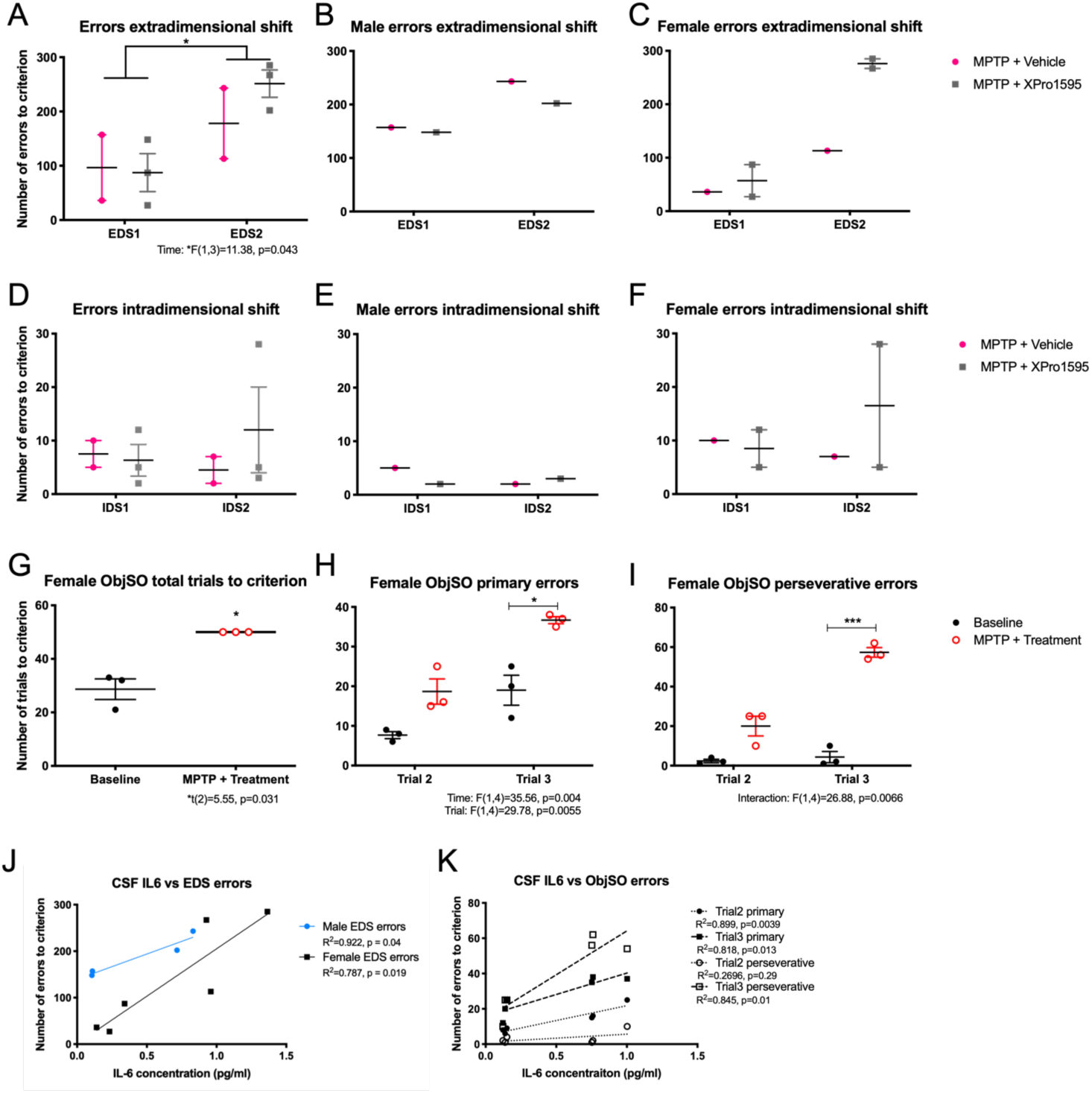
Chronic MPTP impairs cognitive flexibility and significantly increased perseverative behavior. Grouped number of errors during extradimensional shift stages was significantly increased after long-term chronic MPTP treatment (EDS2) compared to short-term MPTP treatment (EDS1;A). Qualitative analysis of the same measurements by sex reveal that females performed EDS1 with less errors than males at the same timepoint, yet both sexes exhibited increased errors following long-term MPTP administration (B,C). Intradimensional shift stage errors did not show any significant differences across the group or within sexes when comparing baseline (IDS1) to late stage chronic MPTP (IDS2; D-F), suggesting intact associative conditioning. In females, object self-order task (ObjSO) results reveal a significant increase in trials to reach criterion post-MPTP compared to baseline where primary and perseverative errors were significantly higher in the 3^rd^ trials (H,I). EDS errors to reach criterion were also correlated with central levels of IL-6 (J). Both errors during trial 3 significantly correlated to animals clinical rating status and CSF IL-6 levels (K).

### MPTP altered gut permeability

Intestinal permeability was evaluated with the use of a cocktail that includes sugars that are passively absorbed from the gut under either transcellular or paracellular mechanisms without considerable metabolism and then excreted into the urine. A two-way ANOVA evaluation of the % of sugar excreted three hours after oral delivery of the sugar solution showed no effect of sex (Lactulose: F(1,3)=0.1616, p=0.7146, mannitol: F(1,3)=0.209, p=0.6786, sucralose: F(1,3)=0.2128, p=0.676, sucrose: F(1,3)=0.5924, p=0.4976) at either baseline or MPTP timepoints. Because there were no detectable differences between the sexes, the monkeys were combined in the analysis to focus the evaluation on the effect of MPTP on intestinal permeability. Monkeys treated with 11 weeks of systemic MPTP displayed no significant effects on gastro-duodenal or small intestinal permeability (**Figure S6A-D**). No significant drug treatment effects were detected in the sugar absorption response; however, a trend towards increases in lactulose (p=0.094) was observed at endpoint compared to early MPTP in both XPro1595- and vehicle-treated animals, suggesting that continued MPTP administration increased permeability in the small intestine (**Figure S6E-F**).

### MPTP influenced fecal microbial diversity and composition with sex differences being demonstrated in gut microbiota and SCFA metabolites

Analyses of alpha diversity indices (variance within-sample) and beta diversity (variance between-samples) were used to examine changes in fecal microbial community structure within each monkey and between experimental conditions. Alpha diversity indices (i.e., Shannon, Simpson, richness, and evenness) were examined at different taxonomic levels (Phylum, class, order, family, genus, and species). Alpha diversity indices indicted that stool samples from monkeys obtained after 11 weeks of low-dose MPTP displayed similar measurements when compared to baseline. However, a significant effect of sex was found at the taxonomic level of phylum Shannon, Simpson, richness and evenness measurements indicating increased diversity in males compared to females when challenged chronically with MPTP (**Figure 8A-D**). Similar significant effects of sex were seen in alpha diversity at other taxonomic levels (class, order, and family; data not shown). After drug treatment, we observed an increase in richness as a % of baseline (i.e., the total number of species per sample), at the taxonomic level of phylum (F(1,3)=15.8, p=0.0285) for monkeys receiving XPro1595 compared to the pattern of reduced richness found in vehicle-treated animals. A similar trend in richness was observed at the taxonomic level of family (F(1,3)=9.385, p=0.0548; **Figure 8E-F**).

**Figure 8.**
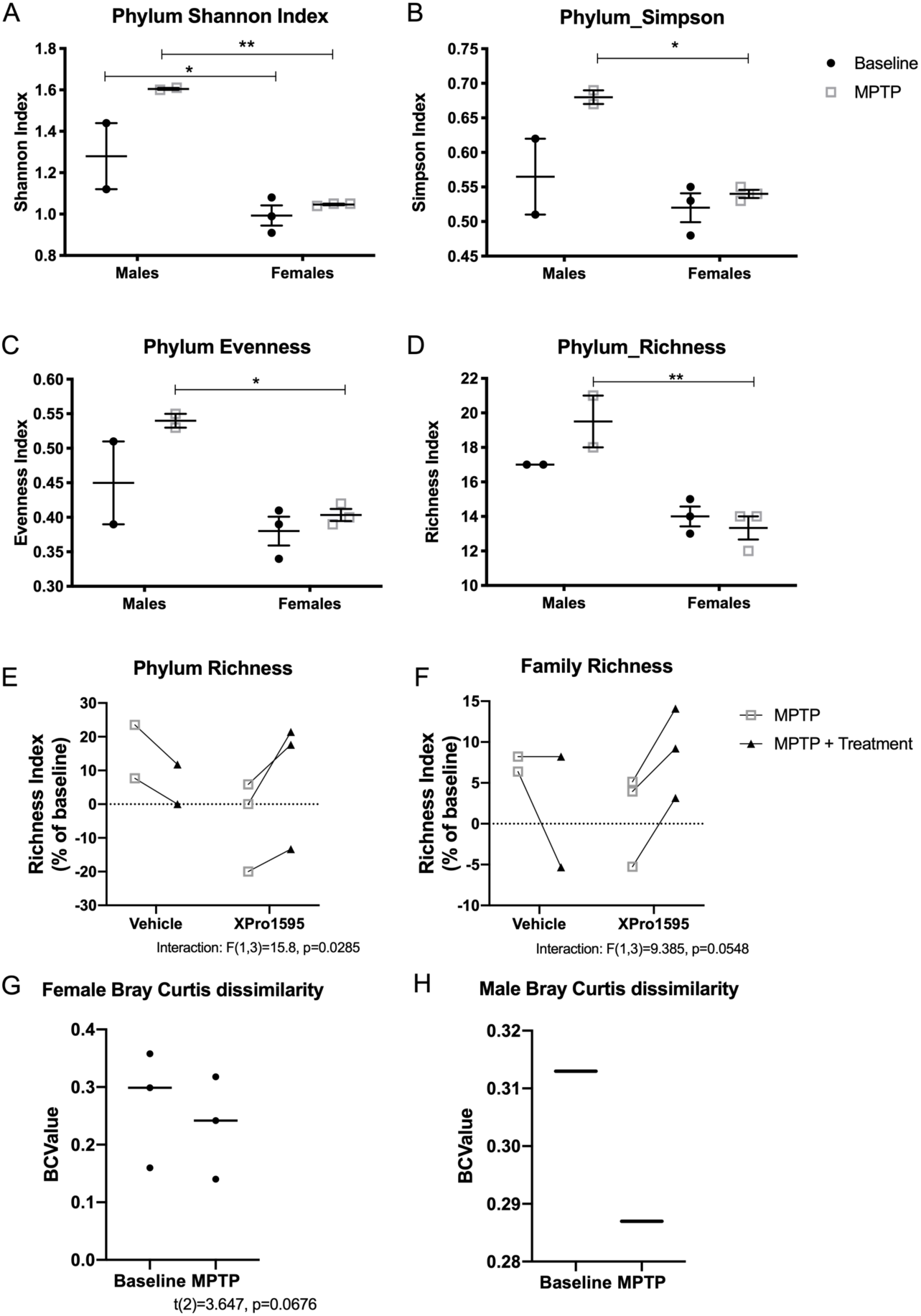
Chronic low-dose MPTP increases alpha diversity in males compared to females. Alpha diversity measures of Shannon, Simpson, Evenness and Richness indices for phylum taxa (A-D), depicted at the phylum taxonomic level, are increased in males compared to females after low doses of systemic MPTP when animals did not present with motor phenotype (Phylum Shannon F(1,3)=38.22, P=0.0085, Phylum Simpson F(1,3)=12.51, p=0.00385, Phylum Evenness F(1,3)=26.21, P=0.0144, Phylum Richness F(1,3)=58.55, P=0.0046). Similar patterns were found in identical alpha diversity indices at lower taxonomic levels including class, order and family (not genus nor species, data not shown). Evaluation of the % change of richness index, at both the phylum and family taxonomic levels (E-F) suggested the total number of bacteria differs in animals treated with XPro1595 compared to vehicle (Phylum Richness interaction F(1,3)=15.8, p=0.0285; Family Richness interaction F(1,3)=9.385, p=0.0548). The baseline genus microbial composition in female display trends towards dissimilarity (p=0.0676), when compared to their corresponding MPTP measures (G). Only trend lines can be compared between male timepoints (n=1; H). Values closest to 1 represent dissimilarity in species populations while values closest to 0 are most similar. MPTP = 10 weeks of MPTP dosing and pre-drug treatment; MPTP + treatment = endpoint measurement post-MPTP and post-drug treatment after 26 weeks of MPTP in males and 40 weeks of MPTP in females.

To further compare microbial community structure, beta diversity (variance between animal samples) was conducted using Bray-Curtis pairwise dissimilarity analyses (0=similarity; 1=dissimilarity) to compare the overall microbial compositional differences between baseline and short-term MPTP administration. The baseline genus microbial composition in females displayed a trend towards dissimilarity but was not statistically significant (Female p=0.0676). This metric was not calculated for males, or for the final effect of treatment, given the limitations of our sample size (**Figure 8G-H**).

Given the impact of PD on colonic bacterial composition reported by multiple groups (Keshavarzian et al., 2015; Scheperjans et al., 2015; Unger et al., 2016), we further evaluated the relative abundance of individual taxa at different taxonomic levels. Visual observations of the mean phylum abundances in both the males and females at either baseline or after 11 weeks of low doses of MPTP revealed minor, yet detectable differences between sexes (Figure 9A). A reduction in the relative abundance of phylum Firmicutes, plus the ratio of Firmicutes to Bacteriodetes ratio was evident in males after short-term MPTP treatment (Figure 9B-D). Conversely, the phylum Verrucomicrobia was markedly increased in males treated with MPTP compared to females treated with MPTP (Figure 9D). Similarly, the relative abundance of bacterial RFP12 (within the Verrucomicrobia phylum) was significantly increased in male monkeys treated with MPTP when compared to baseline (p<0.0001; Figure S7). No other significant differences of the effect of MPTP were found at the taxonomic level of family including the commonly evaluated bacterial family Prevotellaceae, which has been reported to be reduced in PD patients (Scheperjans et al., 2015; Unger et al., 2016). At the taxonomic level of genus, the relative abundance profiles demonstrate a striking reduction of highly abundant taxa in males at either timepoints compared to female monkeys (Figure 9E). Two-way analysis of sex and time revealed the average abundance of bacterial genus Blautia, a known beneficial SCFA-producing bacteria, to be significantly decreased in female monkeys after MPTP treatment (Figure 9F). Due to the low sample size and impact of sex on microbial diversity, we did not attempt to evaluate the effect of XPro1595 on taxa abundances.

**Figure 9.**
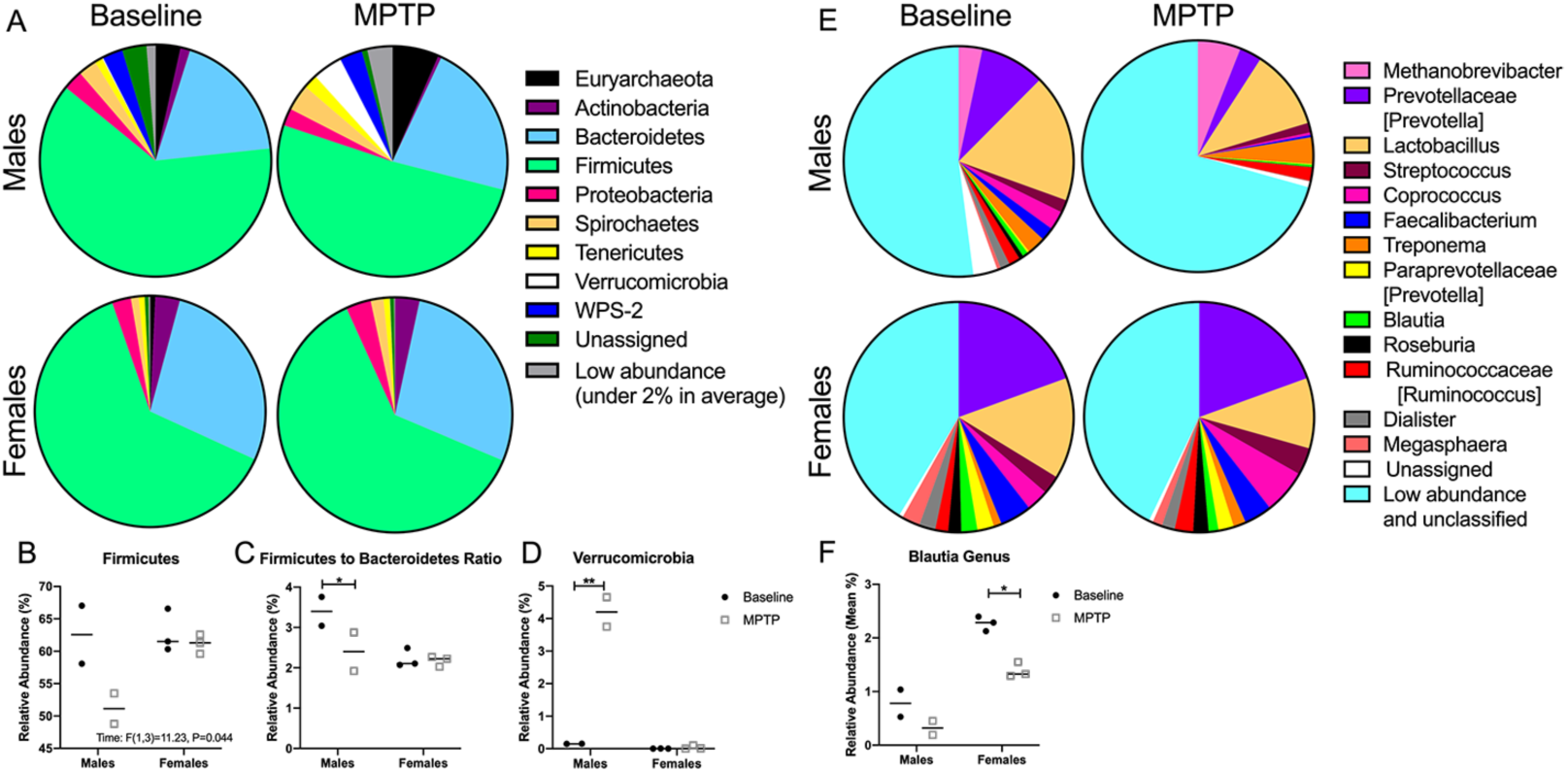
Gut microbiota relative abundance of phylum and genus individual taxa differs between sexes. Pie charts display the average percent relative abundance of bacterial phylum individual taxa in male (n=2) and female (n=3) monkeys at baseline and after 10 weeks of chronic MPTP (A). At the taxonomic level of phylum, MPTP administration has a greater effect in males than females showing a significant reduction in the abundances of *Firmicutes* (B) and *Firmicutes* to *Bacteroidetes* ratio (C) and increased *Verrucomicrobia* (D). Pie charts display the average percent relative abundance of genus individual taxa in male (n=2) and female (n=3) monkeys at baseline and after 11 weeks of chronic MPTP (E). Females display a decreased relative abundance of the bacterial genus Blautia following short-term MPTP dosing (F). Post-hoc analysis was conducted using Sidak’s multiple comparisons. *p<0.05, **<0.01

SCFA are produced from gut bacteria fermentation of dietary carbohydrates and have been shown to inhibit inflammation (MacFabe, 2015). Investigation of SCFA stool levels at baseline and MPTP revealed increased levels of acetate, butyrate, propionate, and total SCFA in female fecal samples compared to males with no significant effect of MPTP (**Figure 10A-D**). Butyrate levels in stool positively correlated with the total number of bacterial family *Lachnospiraceae* sequences, a SCFA-producing bacteria known to produce butyric acid (R^2^=0.808, p=0.038). Following treatment, the % change in SCFA levels compared to baseline displayed upward trends in stool acetate, propionate and total SCFA levels at endpoint following MPTP and drug treatment with no difference between XPro1595 and vehicle-treated monkeys (**Figure 10E-G**). Sex difference were also seen in lipopolysaccharide binding protein (LBP) serum levels such that baseline levels were elevated in males compared to female monkeys, and males had a significant reduction in LBP levels after 10 weeks of MPTP dosing with no changes noted in female serum (**Figure 11**). No significant change was found in LPS or LBP secretion as a % change from baseline between XPro1595- and vehicle-treated animals (LPS: F(1,3)=0.83, p=0.429; LBP: F(1,3)=1.28, p=0.339; data not shown), therefore analysis was conducted with aniamls grouped by sex. Comparison of LPS and LBP (as % from baseline) at 10 weeks of MPTP (pre-drug treatment) dosing and endpoint (post-drug treatment) or 26 or 40 weeks of MPTP for males and females, respectively, demonstrates significant increases in the circulating levels of LPS and LBP in males compared to females (**Figure 11C,D**).

**Figure 10.**
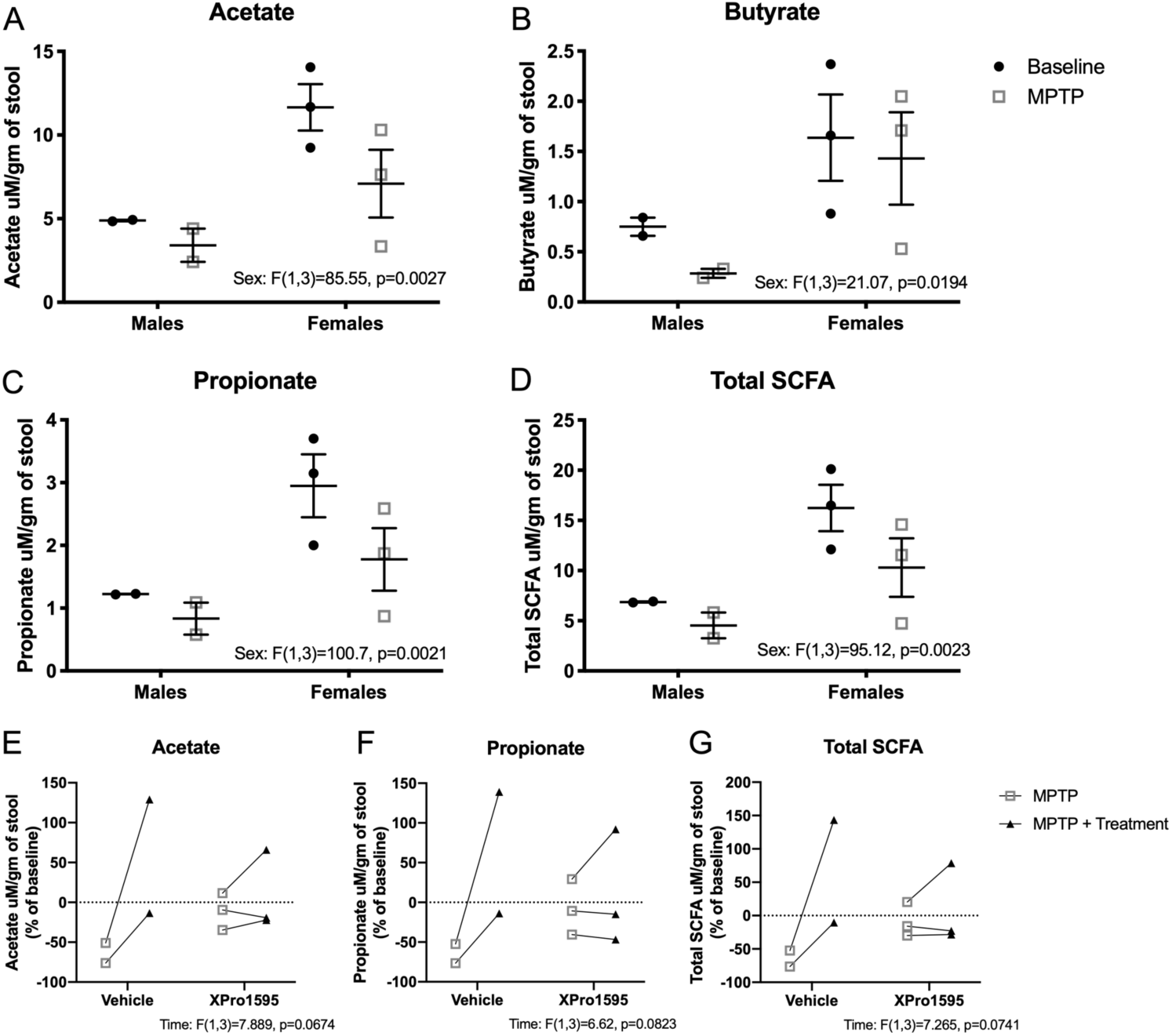
Female monkeys have increased levels of short chain fatty acids in stool at baseline. Two-way ANOVA analysis revealed SCFA acetate, butyrate, propionate and total SCFA levels were significantly increased in female monkeys (A-D). Acetate (E), propionate (F) and total SCFA (G) display trends towards an increase at endpoint following MPTP and drug treatment with no difference between XPro1595 and vehicle-treated monkeys. MPTP = 10 weeks of MPTP dosing and pre-drug treatment; MPTP + treatment = endpoint measurement post-MPTP and post-drug treatment after 26 weeks of MPTP in males and 40 weeks of MPTP in females.

**Figure 11.**
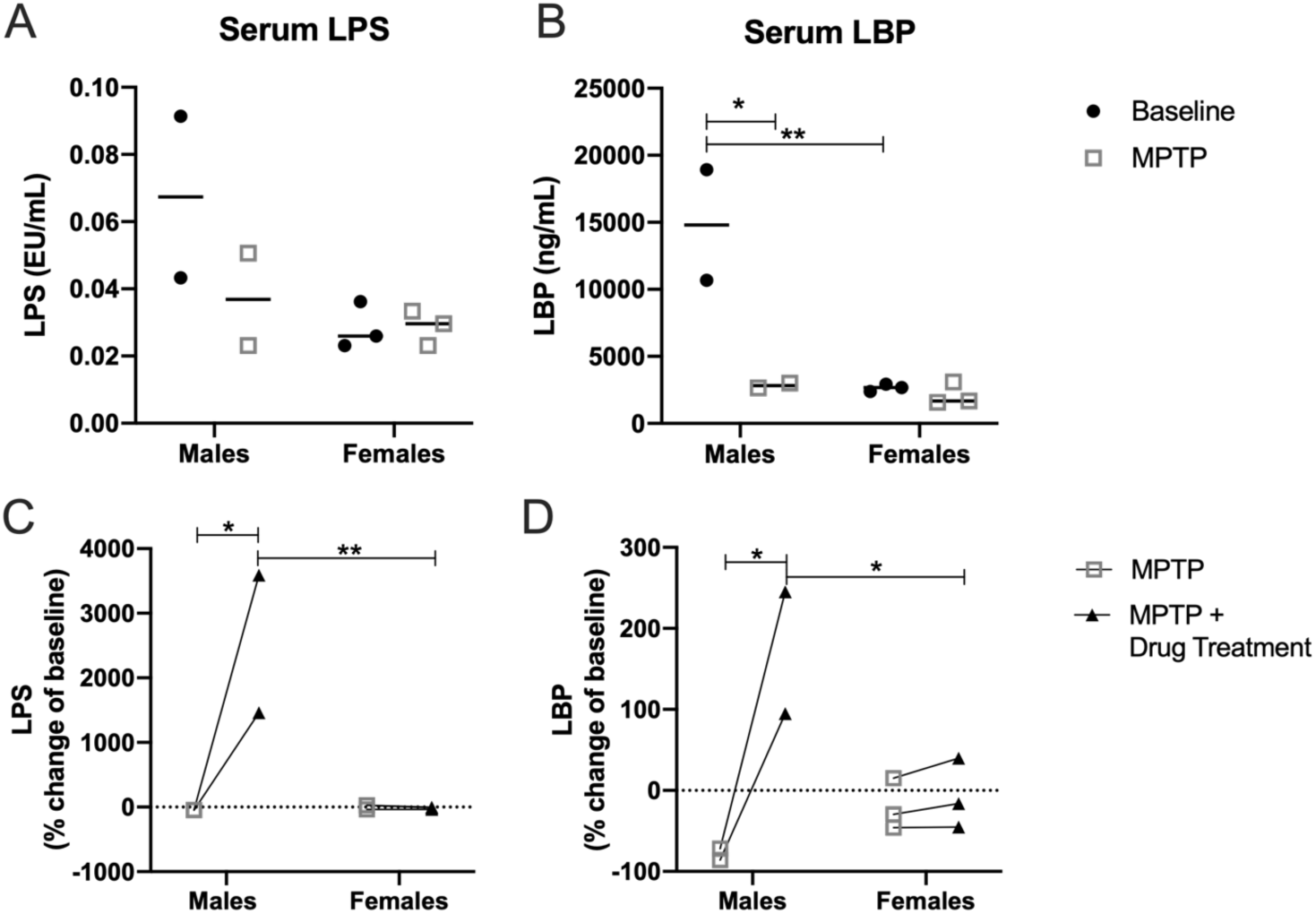
Male monkeys display higher baseline serum LPS and LBP than females and chronic MPTP dosing exacerbates this difference. Serum levels of lipopolysaccharide (LPS, A) and lipopolysaccharide binding protein (LBP, B) reveal male monkeys have significantly higher serum LBP baseline levels and demonstrate a large drop in levels in response to 10 weeks of MPTP. Following drug treatment and escalating MPTP injections, repeated MPTP injections to males reveal greater increases in LPS and LBP as a % of baseline compared to females (C,D). *p<0.05, **p<0.01

Anatomical sections of the gastrointestinal tract were evaluated postmortem for the inflammatory markers CD68 and HLA-DR. Within the lower colon, crypts presented significantly less CD68 immunoreactivity (CD68-ir) by optical intensity and % area in XPro1595-compared to vehicle-treated animals (**Figure 12**). However, colorectal biopsies did not display any significant changes in cytokines IL1β, IL-6 or IL-2 following drug treatment, yet a trend was found for increased IL-6 with escalating delivery of MPTP (**Figure S8**). No effect of sex was observed for CD68 quantification in the gut. Additionally, no differences were found in HLA-DR-ir in either ileum or lower colon (data not shown).

**Figure 12.**
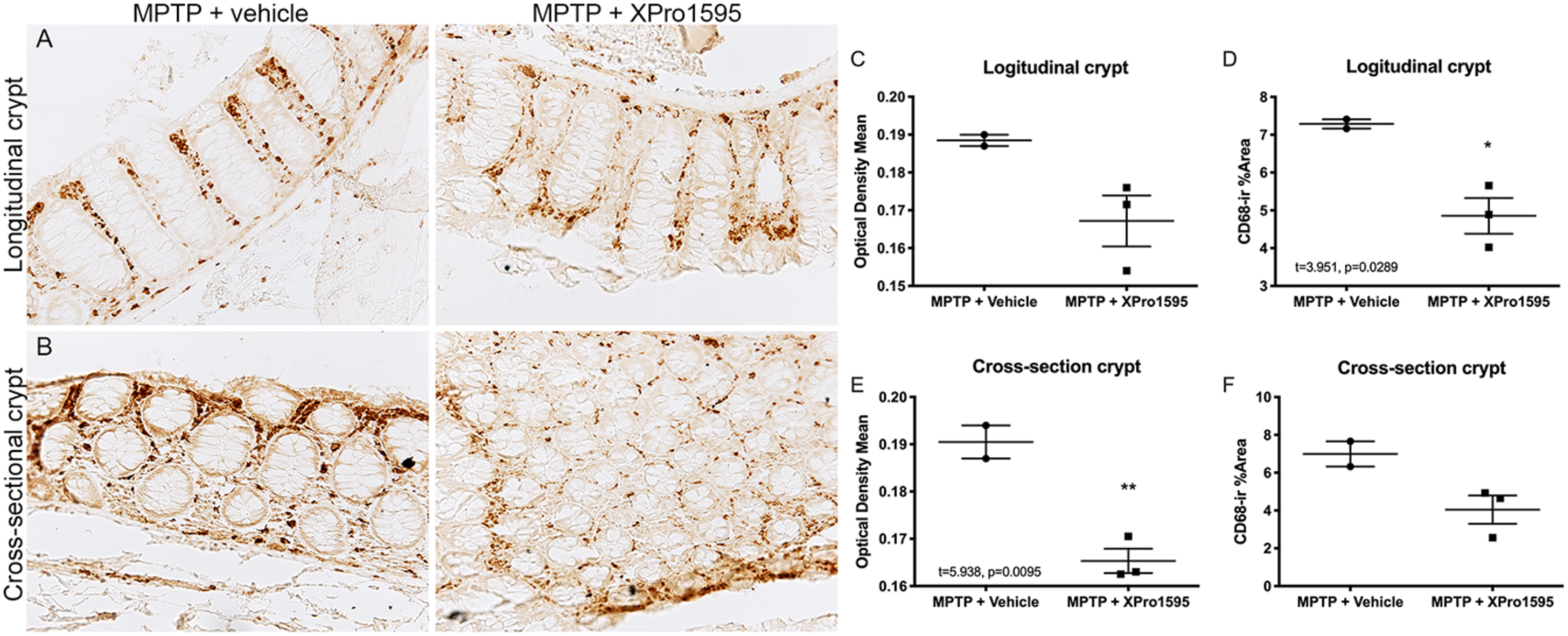
Treatment with XPro1595 reduced CD68-ir in the lower colon. Representative microphotographs of CD68 staining in the crypts of the lower colon either in longitudinal (A) or cross-sectional orientation (B). Quantification of CD68-ir optical density (C,E) and % area coverage (D,F) demonstrate significant reductions in the colon of XPro1595-treated monkeys compared to vehicle-treated.

## 4. Discussion

The overwhelming amount of evidence implicating a role for neuroinflammation in PD pathophysiology now compels our field to enrich for and enroll PD subjects for immunomodulatory clinical trials to better identify prognostic biomarkers, robust outcome measures, and therapeutic interventions. This is especially critical as various inflammatory targets become validated and/or drugs shown to be beneficial in autoimmune diseases are repurposed for use in the PD clinic. While a multitude of studies in rodents including several from our group suggest neuroinflammation starts early, precedes degeneration of the nigrostriatal pathway and, if blocked early, could mitigate degeneration of vulnerable populations, there is valid concern that results from those studies may have limited translational value to humans due to immunological differences between species; therefore, studies in non-human primates are merited to improve the probability that findings will translate from the lab to the clinic. Here we report the first attempt to evaluate the impact of progressive parkinsonism (modeled by chronic low doses of MPTP) on central and peripheral inflammatory responses in rhesus monkeys. Additionally, we investigated nonmotor symptoms that arise in this model including the first assessment of microbiome in MPTP monkeys, and the extent to which a sTNF-specific biologic XPro1595 (presently in clinical trials in early Alzheimer’s disease) could mitigate inflammatory responses and ameliorate nonmotor symptoms.

### Sex-specific immune responses to MPTP

Our novel results suggest that non-human primates display striking sex-specific differences in neuroinflammatory responses to a neurotoxic insult delivered systemically that elicits chronic progressive degeneration of dopaminergic and non-dopaminergic cell groups. Specifically, MPTP-induced inflammation and microglial activation as measured by circulating levels of inflammatory markers and [^18^F]FEPPA PET respectively was greater and triggered earlier in male than in female macaques. In female monkeys, no significant increase in [^18^F]FEPPA binding was detectable at any neuroimaging time point and NGAL and CRP levels measured in biofluids did not increase during the course of the chronic MPTP delivery. Additionally, CSF levels of IL-6 and IFNγ and plasma levels of IL-8 significantly increased over time regardless of sex, suggesting their potential use as a biomarker of MPTP-induced parkinsonism. There have been previous reports of heightened circulating cytokine levels in MPTP-treated monkeys. Specifically, levels of circulating TNF and IFNγ were still increased in male macaques when measured at least one year after MPTP administration, with highest levels in severely parkinsonian animals (Barcia et al., 2005; Barcia et al., 2011). Very few female monkeys were represented in these studies and cytokine analysis was not evaluated for sex effects. In another study, a MPTP-induced increase in TSPO was detected in the male baboon brain using [^11^C]PK11195 PET (Chen et al., 2008). Interestingly, the pattern of [^11^C]PK11195 binding paralleled that found in the males in this study with an early significant increase and gradual decline to near baseline levels. The early increase in TSPO measured by PET scans prior to the beginning of XPro1595 treatment to inhibit sTNF indicates that the window for therapeutic effects derived from sTNF neutralization was missed in these animals. On the other hand, the lack of robust PET signal in female monkeys after MPTP administration prevented us from assessing the effects of sTNF neutralization on global brain inflammation using PET. Although these observations must be confirmed in larger cohorts of animals, they suggest that sex is an important factor to consider in the trial design of future immunomodulatory intervention trials (Barnum et al., 2014).

There is a sexual dimorphic bias in the incidence of PD, such that it presents more in men than women with an age of onset 2 years earlier in men (Twelves et al., 2003). This feature has been recognized in preclinical models showing that male mice develop more severe MPTP-induced neurotoxicity than females (Antzoulatos et al., 2010; Przedborski et al., 2001). However, the mechanisms underlying the development of PD regulated by sex remain unclear and merit further investigation. Several studies have identified an influence of sex on the inflammatory state of PD patients (Houser et al., 2018; Manocha et al., 2017). Additionally, leucine-rich repeated kinase-2 (LRRK2) mutations, which account for 3% of sporadic PD cases and have been associated with dysregulation of the immune system, have been found to bias toward female patients in multiple cohorts (Cilia et al., 2014; Gan-Or et al., 2015; Marder et al., 2015), highlighting the complex effects of sex and immunity on disease pathogenesis. There are also direct and indirect actions of immunomodulation of sex hormones that may influence disease. Estrogen has also been linked to anti-inflammatory activity in the brain, such that it dampens microglial activation induced by lipopolysaccharide exposure (Vegeto et al., 2008; Vegeto et al., 2001). Furthermore, striatal dopamine depletion induced by MPTP was mitigated following estradiol hormone replacement in ovariectomized mice and monkeys (Liu and Dluzen, 2007; Miller et al., 1998; Morissette and Di Paolo, 2009), suggesting that estrogen may exert a neuroprotective role. Interestingly, women often have less severe motor deficits than men in advanced PD stages and respond better to levodopa therapy (Horstink et al., 2007; Zappia et al., 2005). Because estrogen influences multiple systems, the mechanism of its neuroprotective properties is not well understood. The monkeys used in this study are considered adults and although we did not measure circulating hormones, we expect the females to have had normal estrous cycles and this may have contributed to the sex-specific responses to chronic administration of MPTP.

Enhanced neuroinflammation is critical in promoting neuronal damage; therefore, a robust and early inflammatory response to MPTP that does not resolve would be expected to contribute to a faster rate of disease progression in males. Additionally, we show a marked divergence between sexes in the levels of NGAL CSF and plasma (**Figure 5F,G**) and CCL2 CSF (F(1,3)=9.208, p=0.05; data not shown) over time, such that males have intrinsically higher levels of these factors in their biofluids. NGAL (or lipocalin-2) has shown to heighten the sensitivity of neurons to amyloid beta toxicity in murine primary cortical cultures (Naude et al., 2012). While, CCL2 signaling recruits peripheral immune cells into the brain, and the crossing of peripheral monocytes has shown to be detrimental as demonstrated in animals models of multiple sclerosis and PD (Ajami et al., 2011; Harms et al., 2018); however, their exact role in the CNS is not clear. Taken together, our results suggest that increased basal levels of NGAL and CCL2 may influence the heighted male vulnerability to MPTP toxicity. Interestingly, although males were administered less total MPTP than females due to their shortened timeline, a similar level of nigrostriatal dopamine neurodegeneration was found at endpoint in both males and females, as measured by stereological TH+ cell counts in the SN and quantification of the final striatal and nigral [^18^F]FECNT binding, suggesting that the sensitivity of the nigrostriatal system to neuroinflammatory stress is different between the sexes. However, it is still not clear if the divergent progression of motor impairment and nigrostriatal degeneration is attributed to the sex differences found in basal levels of immune mediators, immune responses to MPTP or to an interaction between estrogen and dopaminergic vulnerability to the neurotoxin.

### Nigral [^18^F]FEPPA binding and IBA1+ surface area in good agreement

Previous studies have found good correlations between TSPO PET or autoradiography binding and histological immunoreactivity of innate immune cells positive for CD68 (Venneti et al., 2007; Vowinckel et al., 1997) and CD11b (Converse et al., 2011; Dickens et al., 2014). Our results indicate that IBA1 surface area is a good surrogate measurement for *in vivo* [^18^F]FEPPA binding. IBA1 is specific to microglia, and the increased surface area is suggestive of their activation state when TSPO is reported to be elevated. Increased surface area of IBA1 immunofluorescence in females compared to male monkeys corroborates the [^18^F]FEPPA results that demonstrated greater levels of TSPO at endpoint in the females than males. However, when evaluating levels of antigen presentation proteins (HLA-DR) within the IBA1+ microglia, we found no significant relationship with [^18^F]FEPPA binding; but that is not surprising as not all microglia effector functions are likely to be reflected in the activation status measured by upregulation of TPSO protein detected by [^18^F]FEPPA. While there are significant limitations to the use of TSPO ligands as specific indices of microglia activation, obtaining consistent outcomes from independent neuroimaging and neurohistological approaches give us additional confidence in the findings.

### Monkeys develop cognitive dysfunction following chronic MPTP

MPTP-treated monkey models have been widely used to study the clinical motor symptoms of PD because of the severe degeneration of the nigrostriatal dopaminergic system. However, based on evidence that chronic systemic MPTP administration also induces loss of noradrenergic, serotonergic and intralaminar thalamic neurons, this particular model is also useful for studying PD-like nonmotor symptoms including cognitive deficits (McDowell and Chesselet, 2012). In this study, both male and female monkeys presented with deficits in attention set-shifting prior to significant motor impairment. The EDS2 trials began 17 weeks after the start of MPTP administration and continued for up to 21 weeks depending on the animal’s performance ability, and the animal’s number of errors significantly increased during this task. Yet, after 22-23 weeks of MPTP treatment, 4 of the 5 animals successfully performed the IDS tasks, demonstrating that MPTP did not affect associative learning and instead had specific effects on complex cognitive flexibility to shift attention to an external dimension. The CRS of male monkeys did not significantly increase until week 23 of MPTP treatment, while 30 weeks of MPTP intoxication were needed to induce motor impairments in female monkeys. The order of behavior appearance suggests that low chronic intramuscular doses of MPTP are able to model the early cognitive dysfunction that has become more recognized in PD patients (Elgh et al., 2009; Muslimovic et al., 2005). Additionally, the increased primary and perseverative errors found in the third trial of the ObjSO task, which was performed between 30 and 35 weeks of MPTP, suggest that working memory is impaired in female monkeys following chronic long-term exposure to MPTP. Our data extend results of previous studies reporting deficits in attention and executive function in rhesus monkeys given low doses of intravenous MPTP (Decamp and Schneider, 2004). Associations have been reported between cognitive decline and inflammatory cytokines (Menza et al., 2010). Monkeys of both sexes demonstrated increases in IL-6 in CSF and plasma and in IFNγ in CSF prior to their measurable cognitive deficits. We were not able to determine whether blockade of sTNF had an impact on cognitive symptoms, due to the low number of animals and sex-specific response to chronic MPTP administration. Additional studies will be needed to determine if neutralization of sTNF with XPro1595 is protective against cognitive impairments in male and female MPTP-treated monkeys.

### MPTP and sex-specific responses in gastrointestinal measures

Considerable interest has been given to the gut-brain connection in PD with multiple studies investigating the role of microbiome and gut permeability in disease pathophysiology in humans and animal models. To our knowledge, this study is the first to evaluate intestinal permeability, microbiota and targeted SCFA metabolomics from MPTP-treated monkeys. Previous studies in patients with PD have reported hyperpermeable intestinal epithelium as evinced by increased excreted sucralose 24 hours following ingestion of an oral sugar solution and decreased plasma LBP (Forsyth et al., 2011). Forsyth et al also reported that 5-hour urinary lactulose and mannitol were normal in these PD patients, suggesting that disruption of intestinal brier integrity (“leaky gut”) in early PD is primarily in the colon. In this study, it was not possible to collect urine for 24 hours; and therefore, we could not assess colonic permeability. In addition, we did not find any significant leakiness of small intestine in the MPTP-treated monkeys during the early phases of parkinsonism as there was no significant changes in 3 hour urinary lactulose or mannitol 10 weeks after start of neurotoxin; but monkeys showed a trend for leaky small intestine at endpoint (MPTP + drug treatment) with increases in lactulose detected 3 hours after oral sugar solution compared to after 11 weeks of MPTP. Lactulose is a disaccharide with a larger structure than mannitol, thereby restricting its permeation to larger pores in the crypts, and increases in excretion within 3 hours are consistent with increased small intestine permeability (Camilleri et al., 2010). Decreased levels of serum LBP were also detected in male monkeys after 11 weeks of low doses of MPTP treatment, while at the final sample collection male monkeys specifically demonstrated heightened levels of both LPS and LBP compared to minor changes observed in female serum levels. These results suggest that increased intestinal permeability in males (but not in females) may have resulted in bacterial translocation into the circulation. Indeed, multiple clinical studies have identified gut microbial dysbiosis in PD (Hasegawa et al., 2015; Keshavarzian et al., 2015; Scheperjans et al., 2015), accompanied by alterations in bacterial metabolites or SCFA profiles (Unger et al., 2016).

In this study, we report significant sex-specific alterations in the relative abundance of different intestinal fecal bacteria in response to MPTP. The relative abundances of phylum *Firmicutes* and the *Firmicutes* to *Bacteroidetes* ratio were both decreased, while the abundance of phylum *Verrucomicrobia* abundance was increased in male monkeys during the prodromal stage (after 11 weeks of low-dose MPTP and prior to motor deficits). Similarly, alterations were found in fecal microbiota profiles of PD patients (Keshavarzian et al., 2015). However, *Blautia*, an anti-inflammatory SCFA-butyrate producing bacteria, was significantly less abundant during prodromal stages in female monkeys compared to their baseline levels, which has also been reported in PD patients (Keshavarzian et al., 2015). Furthermore, the alpha-diversity indices, at multiple taxonomic levels, were observed to be significantly increased in male compared to female monkeys in the early stages of MPTP treatment. Although fecal samples from monkeys treated with MPTP did not reveal changes in the SCFA metabolite concentrations as seen in PD (Unger et al., 2016), there was a notable difference in the basal concentrations of all SCFA metabolites between sexes. Interestingly, females had remarkably increased levels of certain SCFAs with known anti-inflammatory properties (Li et al., 2018). SCFA have been associated with intestinal barrier function and endotoxin translocation to the circulation, however no correlations were found between these measures in this study (Maciejewska et al., 2018). Yet due to the differences in clinical presentations between sexes, our small study precludes conclusions on the causal relationships between gastrointestinal measurements and motor symptoms. However, other studies have highlighted the role of gut dysbiosis in the hallmark motor deficits of PD (Sampson et al., 2016). Results from this study warrant further investigation into the evaluation of sex-specific effects on gut dysbiosis in PD, specifically during early prodromal stages of the disease.

### XPro1595 dampened inflammatory responses and potentially altered microglial phenotype

Our findings demonstrate that, independent of sex, XPro1595 significantly dampened inflammatory responses of IL-6 in CSF and plasma and CCL2 in plasma compared to vehicle treatment. We interpret these findings as evidence that neutralization of sTNF with XPro1595 reduced the ongoing inflammatory processes, regardless of the stage of motor symptoms. We previously reported the neuroprotective effects of XPro1595 in a rat model of PD when drug dosing began 3 days after an intrastriatal injection of 6-OHDA. However, delaying treatment to 14 days after the dopaminergic insult also mitigated inflammation but did not result in neuroprotection, suggesting that early blockade of neuroinflammation is required for neuroprotection of nigral dopaminergic neurons in this rodent model of parkinsonism (Barnum et al., 2014). Regardless of the start of treatment, XPro1595 attenuated neuroinflammation as measured by reduced IBA1+ microglia and GFAP+ astrocytes. We found a similar trend in this study with both IBA1 volume and surface area reduced in the male monkey treated with XPro1595. In the rat study, it was thought that XPro1595 reduced microgliosis, while in this study, when the window of treatment was missed, the morphology of IBA1+ microglia indicate reduced processes and perhaps suggestive of ameboid shaped microglia. This interpretation is in line with the increased HLA-DR found in the same male monkey.

Interestingly, we observed increased expression of HLA-DR in microglia of the male monkey that received XPro1595 but increased CD68+ microglia in females that received XPro1595, suggesting possible differences in MPTP-induced microglial phenotypes in response to inhibition of sTNF. We speculate that perhaps females are able to regulate the inflammatory response that contributed to neurodegeneration more effectively than males, consistent with a slower progression of DA loss and motor symptoms. Further investigations into the biological basis of sex-dependent differences in microglia phenotype in response to chronic MPTP or other neurotoxicants are needed. Differences in microglia morphology between males and females have been reported during development (Lenz et al., 2013; Schwarz et al., 2012). Bilbo and colleagues revealed that female mice had a greater number of microglia and more extensive morphology than males at P60 (Schwarz et al., 2012). Given the sex-specific influence on microglia during development, it is not surprising that they would have distinct responses to stimuli in adulthood.

## 5. Conclusion

This novel and detailed study revealed sex-dependent sensitivity to MPTP that resulted in earlier microglial activation by PET, acute plasma IL-6 and CSF TNF, and earlier parkinsonism as measured by motor deficits in males compared to female monkeys. Sex differences were also identified in microbiota and their targeted SCFA metabolites at both basal levels and in response to chronic MPTP. Even though our data demonstrated that XPro1595 may have had an effect on inflammation measures in the biofluids and reduced CD68 expression in the colon, it was not able to counteract the escalating-dose delivery of MPTP and failed to protect against dopaminergic nigrostriatal loss. However, this conclusion is based on a low number of animals that displayed significantly different experimental timelines due to the sex differences in MPTP sensitivity. The escalating and weekly administration of MPTP in this monkey model may not be ideal when assessing neuroprotective properties of immunomodulatory therapies because of the compounded oxidative toxic effect overtime (in particular increased ROS and decreased mitochondrial function). Non-human primates are valuable experimental models because of their genetic proximity to humans, the close resemblance behaviorally to humans, and microglia heterogeneity with gene expression patterns similar to humans (Geirsdottir et al., 2019). Additional studies with larger group sizes of both sexes would enable confirmation and extension of these novel findings. Nevertheless, our results lay the foundation for the exciting possibility that inflammatory responses to neurotoxic stimuli are sexually dimorphic and should be taken into consideration when initiating or assessing therapeutic immunomodulatory interventions in the clinic.

## Acknowledgements

We thank the Tansey, Smith, and Wichmann labs for useful discussions and Francisco Alvarez, PhD for the use of his lab equipment and providing advice on microglia analysis with Imaris. This work was supported in part by the Parkinson’s Disease Foundation, the Michael J. Fox Foundation for Parkinson’s Research, and the Emory Multiplexed Immunoassay Core (EMIC), which is subsidized by the Emory University School of Medicine and is one of the Emory Integrated Core Facilities. Additional support was provided by the National Center for Advancing Translational Sciences of the National Institutes of Health (UL1TR000454), Yerkes National Primate Center NIH base grant (P51 OD011132) and Graduate and Postdoctoral Training in Environmental Health Science and Toxicology (T32ES012870). The content is solely the responsibility of the authors and does not necessarily reflect the official views of the National Institutes of Health.

## Author contributions

MGT and YS: Conceptualization, Funding acquisition, Methodology, Writing – review & editing. VJ: Project administration, Investigation, Formal analysis, Data curation, Investigation, Writing – Original Draft. GM: Investigation, Data curation, Writing – review & editing. GYC: Investigation, Writing – review & editing. ARW and JB: Methodology, Writing – review & editing. DK: Investigation, Writing – review & editing. TR: Investigation, Writing – review & editing. JN: Methodology, Data Curation, Writing – review & editing. RV and MG: Methodology, Writing – review & editing. LH: Supervision, Writing – review & editing. SG, AN, MS and PE: Investigation, Writing – review & editing. AK: Supervision, Writing – review & editing.

## Supplemental figures (see attached supplemental document)

**Figure S1. Injections of peripheral lipopolysaccharide (LPS) increases global [18F]FEPPA binding in the brain.** (A) Acute and chronic LPS-dosing regimens were performed in two separate challenges separated by a 3-week washout. Two female monkeys received a PET scan using [18F]FEPPA after the washout period (baseline) and again 2 hrs post-acute or chronic LPS dosing. Acute animal (RUM7) received a single dose of 3E4 EU/kg LPS, while the chronically-treated monkey (RYJ10) received incremental doses (3E4, 4.5E4, 6E4 EU/kg) of LPS on Days 0, 3, and 6. Blood was collected at baseline (day 0) immediately prior to intravenous administration of LPS, 2 hrs post-LPS injection, and at end of the PET scan or approximately 4.5 hrs post-LPS injection. (B) Plasma levels of IL-6 and TNF increased after LPS injection(s) as measured with MesoScale multiplexed immunoassays. (C) PET images of [18F]FEPPA binding (SUV) at coronal and sagittal brain levels matched to representative MRI section in acute and chronic LPS animals at baseline and post-LPS. (D) Grouped baseline [18F]FEPPA SUV (mean ±SEM) amongst cortical and subcortical ROIs demonstrate no difference in binding by region. Individual [18F]FEPPA binding % change of baseline demonstrate increased binding post-LPS in both LPS dosed animals.

**Figure S2.** [^18^F]**-FECNT PET demonstrates MPTP toxicity.** Coronal images of [^18^F]FECNT PET in the pre-commisural striatum of chronic MPTP-treated rhesus monkeys that received systemic administration of vehicle (A,C) or XPro1595 (B,D) at baseline and 8 (PET I), 16 (PET II), 24/26 (PET III) and 38 weeks (PET IV) after the start of MPTP treatment. The severity of animal’ clinical rating score is inversely correlated with [^18^F]FECNT standard uptake in various basal ganglia nuclei (I). Increased circulating IL-6 was associated with reduced [^18^F]FECNT in the substantia nigra of male monkeys (J, R^2^=0.529, p=0.041). PA, putamen associative. CA, Caudate associative. PM, putamen motor. AC, accumbens. PL, putamen limbic. SN, substantia nigra. **p <0.01, ***<0.001, ****<0.0001

**Figure S3. Chronic MPTP injections impair fine and gross motor movements.** Fine-motor skill task demonstrates increased treat pick up and total pick up time for males (A,C) and females (B,D) with advanced time. Accelerometer collars recorded motion and revealed a deficit in gross motor movement in all males (E) and females (F) with escalating MPTP dosing. Solid line (vs dashed line) represent animals that received XPro1595.

**Figure S4. Chronic MPTP injections induce significant loss of TH+ cells in nigra and noradrenergic lower brainstem nuclei and TH-ir in the striatum.** Representative microphotographs of TH+ immunohistochemical staining in the substantia nigra (A), lower-brainstem noradrenergic cell groups including the locus coeruleus (LC), A7 and A5 (B), and the striatum at the level of the anterior commissure (C). Unbiased stereological estimate of TH+ cells in the ventral and dorsal substantia nigra (SNv and SNd, respectively) and ventral tegmental area (VTA) reveals significantly decreased cell numbers in MPTP + vehicle and MPTP + XPro1595 animals compared to historical controls (D; F(2,15)=133.6, p<0.0001). Stereological estimate of TH+ cell count in the LC show significant differences between MPTP groups and historical controls (E; F(2,15)=13.7, p=0.0004). Optical density of TH-immunoreactivity in the striatum demonstrated significant reduction in multiple functional regions between MPTP groups and controls (F; F(2,20)=87.2, p<0.0001).*p<0.05, **p<0.01, #p<0.001, $p<0.0001 when compared to controls. %p<0.05 when compared to MPTP + vehicle group.

**Figure S5. XPro1595 levels.** Plasma (A) and CSF (B) were both collected monthly and a cortical biopsy was collected at necropsy for measurements of XPro1595 levels using a human TNF-specific immunoassay (MSD). Normalized plasma and CSF levels of XPro1595 positively correlate (C, R^2^=428, p<0.001). Cortical levels of XPro1595 measured from fresh tissue collected directly prior to perfusion (D). Solid lines (vs dashed lines) represent animals that received XPro1595.

**Figure S6. Gastrointestinal permeability assay**. **Figure S6. Gastrointestinal permeability assay.** No significant changes in sugar absorption were observed in the early period of MPTP dosing (A-D). The % of lactulose (E) and mannitol (F) output post-MPTP/pre-drug treatment and post-MPTP/post-drug treatment timepoints suggest no influence of drug treatment on permeability, yet continued MPTP trended to enhance excretion or alter gut leakiness to lactulose.

**Figure S7. Gut microbiota relative abundance of family taxon differs between sexes**. Pie charts display the average relative abundance of bacterial families in female (n=3) and male (n=2) monkeys at baseline and after 10 weeks of chronic MPTP (A). Males present with increased relative abundance of the bacterial family RFP12 following MPTP dosing (B). Post-hoc analysis was conducted using Sidak’s multiple comparisons. ****p<0.0001

**Figure S8. No significant changes in cytokine levels in gut biopsies after MPTP injections and drug treatment.** Colorectal biopsies were collected at baseline, MPTP and at endpoint following MPTP + drug treatment. Biopsies showed no significant effect of sex or MPTP in IL-1β IL-6 or IL-2 cytokine levels (A-C). Analysis of cytokine levels as a % change from baseline suggests no effect of treatment or time on IL1β and IL-2 and a trend toward increased levels of IL-6 (p=0.0665) at endpoint (MPTP + treatment) compared to MPTP alone (D-F). MPTP = 10 weeks of MPTP dosing and pre-drug treatment; MPTP + treatment = endpoint measurement post-MPTP and post-drug treatment after 26 weeks of MPTP in males and 40 weeks of MPTP in females.

