## Supplemental files for "Sexually dimorphic responses to MPTP found in microglia, inflammation and gut microbiota in a progressive monkey model of Parkinson’s disease"

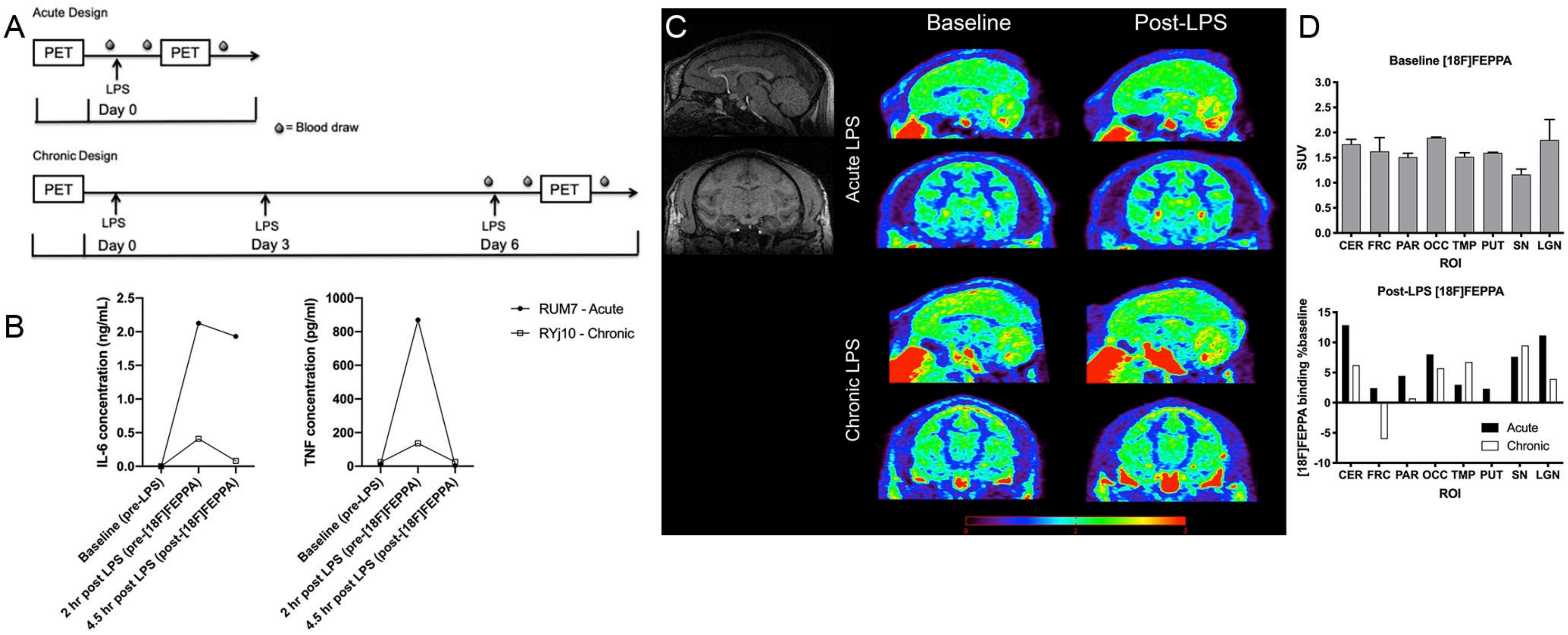


**Figure S1. Injections of peripheral lipopolysaccharide (LPS) increases global [18F]FEPPA binding in the brain.** (A) Acute and chronic LPS-dosing regimens were performed in two separate challenges separated by a 3-week washout. Two female monkeys received a PET scan using [18F]FEPPA after the washout period (baseline) and again 2 hrs post- acute or chronic LPS dosing. Acute animal (RUM7) received a single dose of 3E4 EU/kg LPS, while the chronically-treated monkey (RYJ10) received incremental doses (3E4, 4.5E4, 6E4 EU/kg) of LPS on Days 0, 3, and 6. Blood was collected at baseline (day 0) immediately prior to intravenous administration of LPS, 2 hrs post-LPS injection, and at end of the PET scan or approximately 4.5 hrs post-LPS injection. (B) Plasma levels of IL-6 and TNF increased after LPS injection(s) as measured with MesoScale multiplexed immunoassays. (C) PET images of [18F]FEPPA binding (SUV) at coronal and sagittal brain levels matched to representative MRI section in acute and chronic LPS animals at baseline and post-LPS. (D) Grouped baseline [18F]FEPPA SUV (mean ±SEM) amongst cortical and subcortical ROIs demonstrate no difference in binding by region. Individual [18F]FEPPA binding % change of baseline demonstrate increased binding post-LPS in both LPS dosed animals.

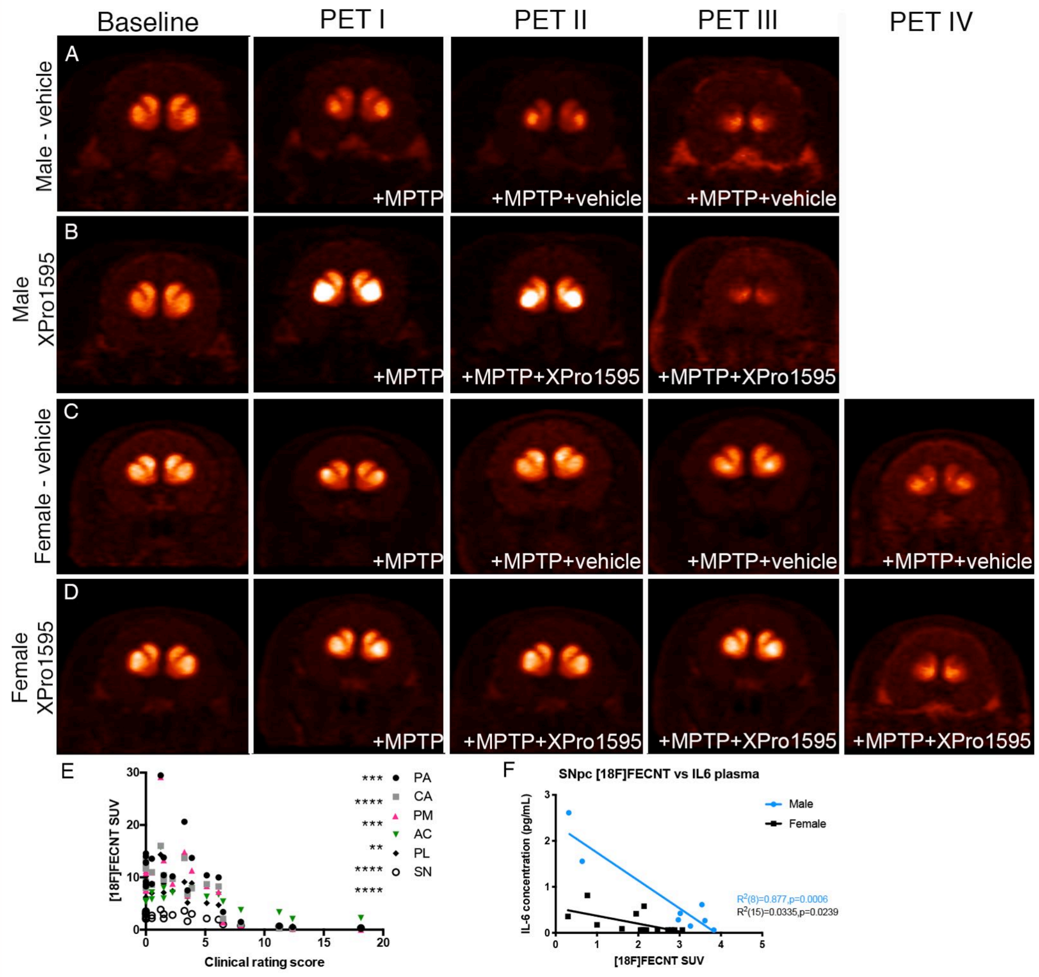


**Figure S2.** [18F]**-FECNT PET demonstrates MPTP toxicity.** Coronal images of [18F]FECNT PET in the pre-commisural striatum of chronic MPTP-treated rhesus monkeys that received systemic administration of vehicle (A,C) or XPro1595 (B,D) at baseline and 8 (PET I), 16 (PET II), 24/26 (PET III) and 38 weeks (PET IV) after the start of MPTP treatment. The severity of animal' clinical rating score is inversely correlated with [18F]FECNT standard uptake in various basal ganglia nuclei (I). Increased circulating IL-6 was associated with reduced [18F]FECNT in the substantia nigra of male monkeys (J, $R^2$=0.529, p=0.041). PA, putamen associative. CA, Caudate associative. PM, putamen motor. AC, accumbens. PL, putamen limbic. SN, substantia nigra. **p <0.01, ***<0.001, ****<0.0001

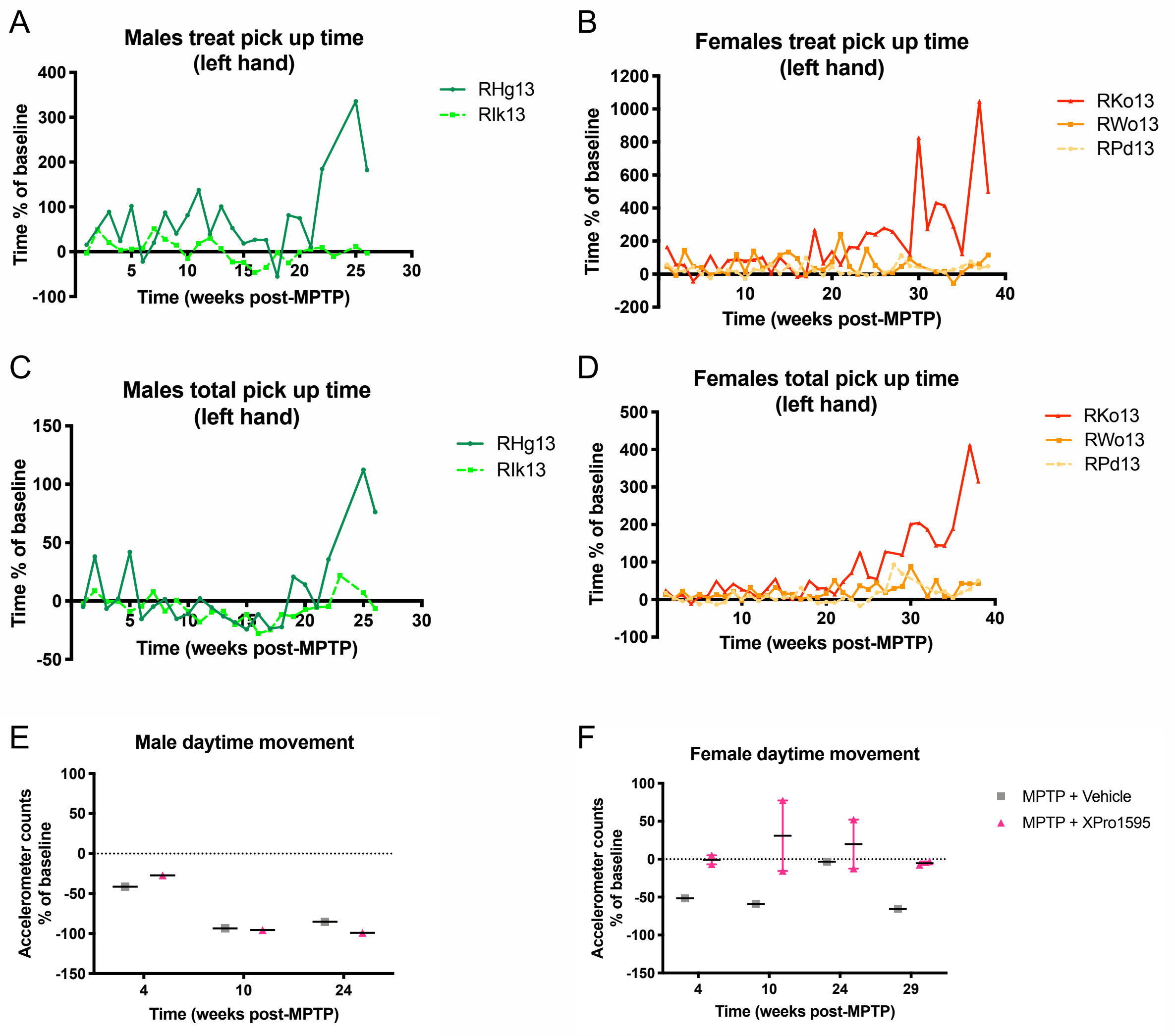


**Figure S3. Chronic MPTP injections impair fine and gross motor movements.** Fine-motor skill task demonstrates increased treat pick up and total pick up time for males (A,C) and females (B,D) with advanced time. Accelerometer collars recorded motion and revealed a deficit in gross motor movement in all males (E) and females (F) with escalating MPTP dosing. Solid line (vs dashed line) represent animals that received XPro1595.

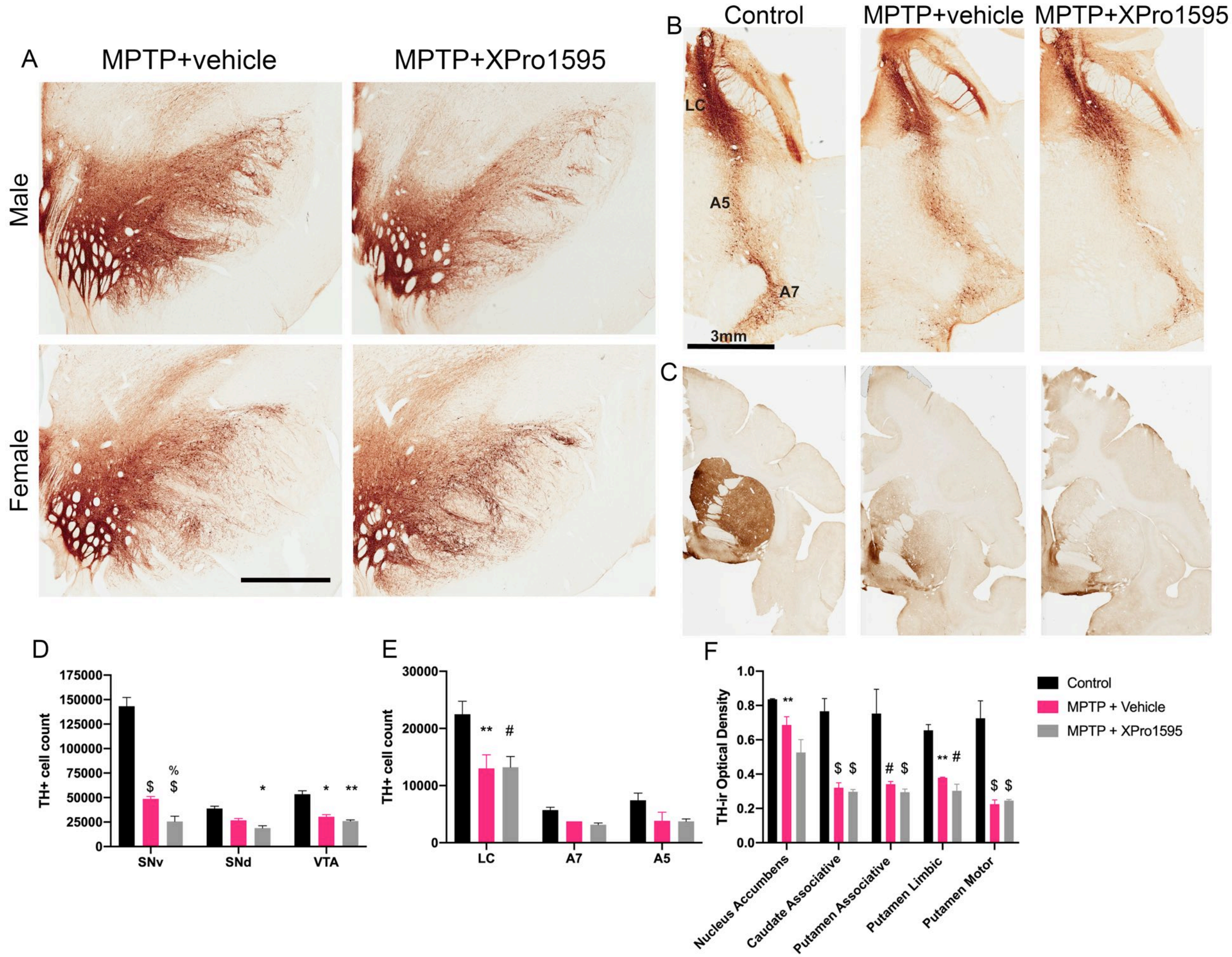


**Figure S4. Chronic MPTP injections induce significant loss of TH+ cells in nigra and noradrenergic lower brainstem nuclei and TH-ir in the striatum.** Representative microphotographs of TH+ immunohistochemical staining in the substantia nigra (A), lower-brainstem noradrenergic cell groups including the locus coeruleus (LC), A7 and A5 (B), and the striatum at the level of the anterior commissure (C). Unbiased stereological estimate of TH+ cells in the ventral and dorsal substantia nigra (SNv and SNd, respectively) and ventral tegmental area (VTA) reveals significantly decreased cell numbers in MPTP + vehicle and MPTP + XPro1595 animals compared to historical controls (D; $F(2,15)=133.6$, $p<0.0001$). Stereological estimate of TH+ cell count in the LC show significant differences between MPTP groups and historical controls (E; $F(2,15)=13.7$, $p=0.0004$). Optical density of TH-immunoreactivity in the striatum demonstrated significant reduction in multiple functional regions between MPTP groups and controls (F; $F(2,20)=87.2$, $p<0.0001$).*$p<0.05$, **$p<0.01$, #$p<0.001$, $\$p<0.0001$ when compared to controls. %$p<0.05$ when compared to MPTP + vehicle group.

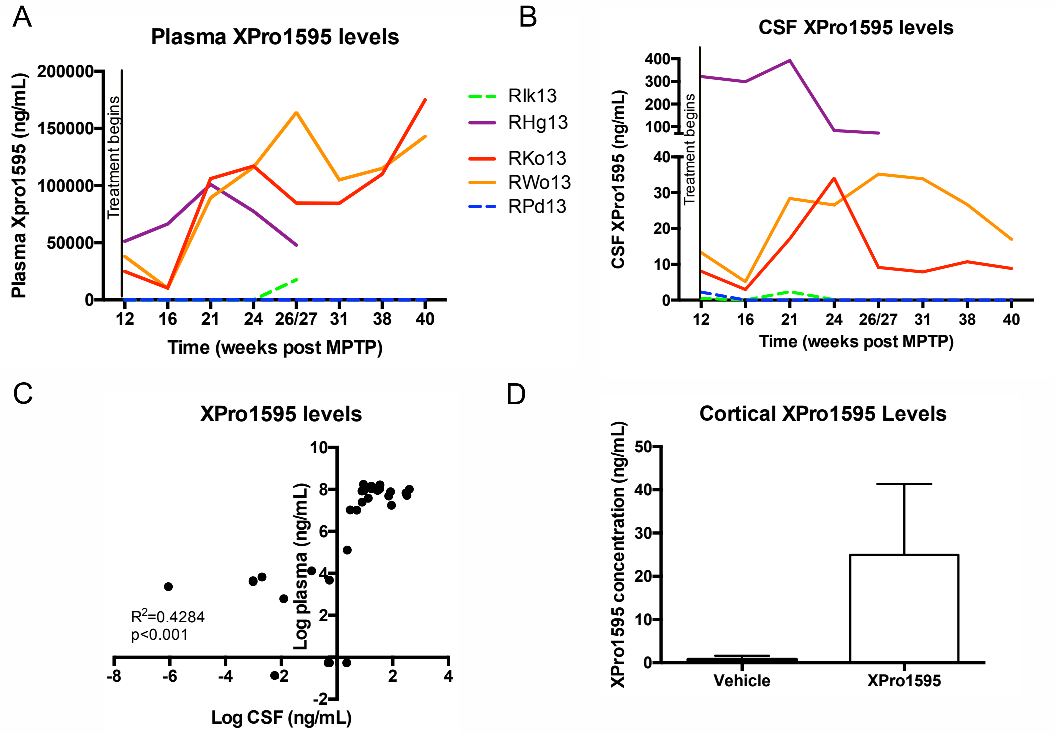


**Figure S5. XPro1595 levels.** Plasma (A) and CSF (B) were both collected monthly and a cortical biopsy was collected at necropsy for measurements of XPro1595 levels using a human TNF-specific immunoassay (MSD). Normalized plasma and CSF levels of XPro1595 positively correlate (C, $R^2$=428, p<0.001). Cortical levels of XPro1595 measured from fresh tissue collected directly prior to perfusion (D). Solid lines (vs dashed lines) represent animals that received XPro1595.

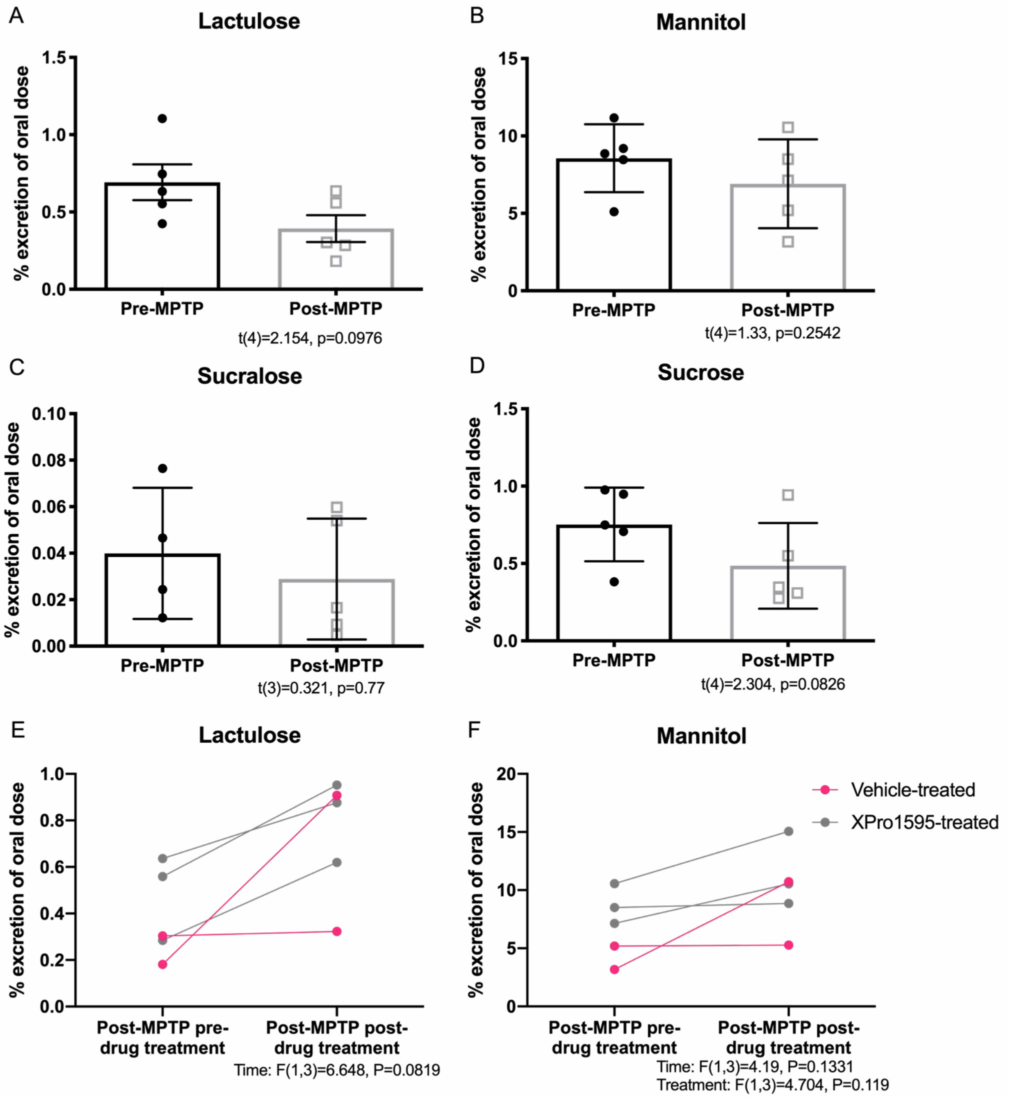


**Figure S6. Gastrointestinal permeability assay. Figure S6. Gastrointestinal permeability assay.** No significant changes in sugar absorption were observed in the early period of MPTP dosing (A-D). The % of lactulose (E) and mannitol (F) output post-MPTP/pre-drug treatment and post-MPTP/post-drug treatment timepoints suggest no influence of drug treatment on permeability, yet continued MPTP trended to enhance excretion or alter gut leakiness to lactulose.

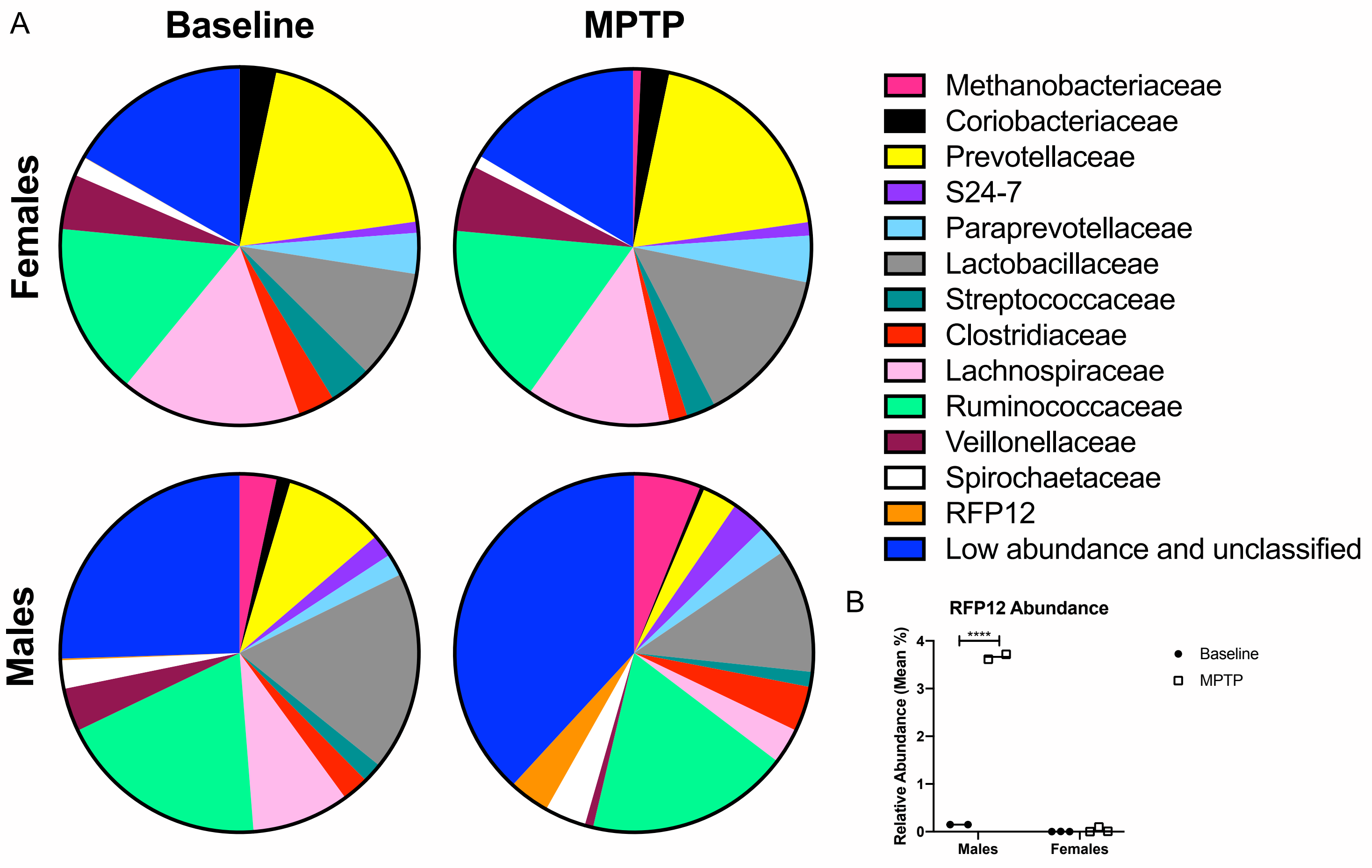


**Figure S7. Gut microbiota relative abundance of family taxon differs between sexes**. Pie charts display the average relative abundance of bacterial families in female (n=3) and male (n=2) monkeys at baseline and after 10 weeks of chronic MPTP (A). Males present with increased relative abundance of the bacterial family RFP12 following MPTP dosing (B). Post-hoc analysis was conducted using Sidak's multiple comparisons. ****$p<0.0001$

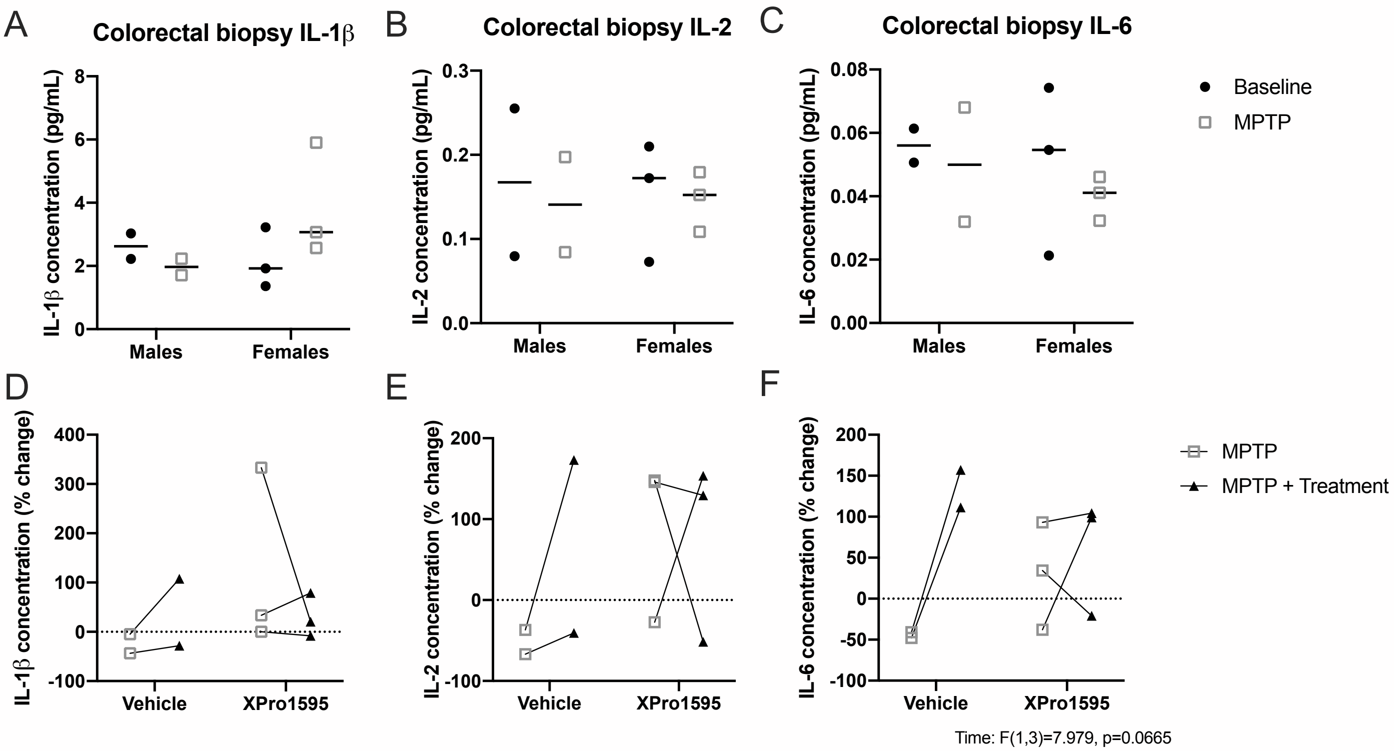


**Figure S8. No significant changes in cytokine levels in gut biopsies after MPTP injections and drug treatment.** Colorectal biopsies were collected at baseline, MPTP and at endpoint following MPTP + drug treatment. Biopsies showed no significant effect of sex or MPTP in IL-1β IL-6 or IL-2 cytokine levels (A-C). Analysis of cytokine levels as a % change from baseline suggests no effect of treatment or time on IL1β and IL-2 and a trend toward increased levels of IL-6 ($p=0.0665$) at endpoint (MPTP + treatment) compared to MPTP alone (D-F). MPTP = 10 weeks of MPTP dosing and pre-drug treatment; MPTP + treatment = endpoint measurement post-MPTP and post-drug treatment after 26 weeks of MPTP in males and 40 weeks of MPTP in females.
